# The invasion pore induced by *Toxoplasma gondii*

**DOI:** 10.1101/2024.10.11.617945

**Authors:** Y. Kegawa, F. Male, I. Jiménez-Munguía, P. S. Blank, E. Mekhedov, G. Ward, J. Zimmerberg

## Abstract

Obligate intracellular parasites invade host cells to survive. Following host cell contact, the apicomplexan *Toxoplasma gondii* injects proteins required for invasion into the host cell. Here, electrophysiological recordings of host cells acquired at sub- 200 ms resolution allowed detection and analysis of a transient increase in host membrane conductance following exposure to *Toxoplasma gondii*. Transients always preceded invasion but parasites depleted of the moving junction protein RON2 generated transients without invading, ruling out a direct structural role for RON2 in generating the conductance pathway or restricting the diffusion of its components. Time-series analysis developed for transients and applied to the entire transient dataset (910,000 data points) revealed multiple quantal conductance changes in the parasite-induced transient, consistent with a rapid insertion, then slower removal, blocking, or inactivation of pore-like conductance steps. Quantal steps for RH had a principal mode with Gaussian mean of 0.26 nS, similar in step size to the apicomplexan protein translocon EXP2. Without RON2 the quantal mean was significantly different (0.19 nS). Because no invasion occurs without poration, the term ‘invasion pore’ is proposed.

## INTRODUCTION

Protozoan pathogens belonging to the phylum Apicomplexa cause life-threatening diseases such as malaria, toxoplasmosis, and cryptosporidiosis in humans and other animals that are important global health burdens with few effective vaccines and limited drugs for treatment (Smith et al., 2021, Striepen, 2013, WHO, 2023). They are obligate intracellular parasites; the intracellular stage of their life cycles begins after host cell invasion. To invade, two types of secretory organelles unique to Apicomplexa are employed: micronemes and rhoptries (Bullen et al., 2019, Carruthers & Sibley, 1997, Dubremetz, 2007, Reviewed in Cova et al., 2022). These organelles contain proteins whose highly coordinated secretion is essential to successful invasion. In *Toxoplasma gondii* (*T. gondii*) tachyzoites, the life cycle stage associated with acute infection, micronemes are abundant organelles mostly located around the parasite’s apical pole (Carruthers & Sibley, 1999). The rhoptries are the largest secretory organelles in the tachyzoite and consist of a rhoptry neck and bulb (Counihan et al., 2013, Proellocks et al., 2010). Between the apical tip of the rhoptry and the parasite plasma membrane lies a small apical vesicle (AV). Current understanding of rhoptry secretion involves fusion of the rhoptry with the AV and fusion of the AV with the parasite plasma membrane, resulting in rhoptry exocytosis (Aquilini et al., 2021, Mageswaran et al., 2021, Sparvoli et al., 2022).

One of the most enigmatic of the molecular events during *T. gondii* invasion is the appearance of rhoptry proteins in the host cell cytoplasm even without complete invasion (aka ‘abortive invasion’) (Håkansson et al., 2001, Ravindran & Boothroyd, 2008, Koshy et al., 2012, Lamarque et al., 2014). The presence of rhoptry proteins in the host cell cytoplasm seems to violate a fundamental cell biological law governing conservation of membrane and secretory protein topology, *i.e.* that protein domains co-translationally inserted into the lumen of the ER remain lumenal throughout the secretory pathway, including within secretory granules and once released from those granules, face the extracellular space. Accordingly, the lumen of the rhoptry is topologically equivalent to the extracellular space and not the host cell cytoplasm. Proteins released from the lumen of the rhoptry can change their topology only by traversing a limiting membrane such as the plasma membrane. The rhoptry neck protein RON2 presents a particularly unique challenge, in that RON2 is a transmembrane protein that is somehow inserted into the host cell plasma membrane after exocytosis, where it serves as a ligand to which AMA1 in the *Toxoplasma* plasma membrane binds and contributes to the formation of a complex of translocated proteins necessary for parasite internalization (the moving junction) (Alexander et al., 2005, Besteiro et al., 2009, Lamarque et al., 2011, Srinivasan et al., 2011, Lamarque et al., 2014). Thus, RON2 is physically proximal to the site of parasite entry.

A transient increase in host cell conductance precedes parasite internalization (Suss-Toby et al., 1996). In a companion paper (Male and Kegawa et al., 2024), this transient increase in host cell ionic permeability (membrane conductance) is also detected as an influx of extracellular Ca^2+^ and is shown to depend on rhoptry exocytosis. Ca^2+^ entry into the host cell occurs at the site of parasite apical contact with the host cell (Male and Kegawa et al., 2024); rhoptry exocytosis also occurs from the apical tip of the parasite. Because the conductance transient and Ca^2+^ entry measure the same event of cell perforation, it is likely that cell perforation occurs at the apical tip, the site of rhoptry protein discharge. Since RON2 is the only known component of the moving junction that is inserted as a transmembrane protein into the host cell plasma membrane, it was reasonable to hypothesize that RON2 contributes to the properties of host cell perforation or that after rhoptry exocytosis RON2-AMA1 binding and formation of the ring-like moving junction at the site of parasite entry might be required to keep the rhoptry secretions localized so as to generate the host cell perforation. Here, we tested these two hypotheses. Neither is correct, because RON2 was not required for either this transient change in conductance to occur or for Ca^2+^ entry. A detailed electrophysiological analysis of the conductance transient revealed the presence of multiple stepwise changes in conductance throughout its trajectory, consistent with a multiple-pore model in the host cell membrane. Based on these results, we propose that these stepwise changes in conductance represent incorporation or formation of discrete membrane structures we term “invasion pores” that may function as a transient breach in the host cell plasma membrane allowing rapid rhoptry protein translocation into the host cytosol.

## RESULTS

### Each invading wild-type T. gondii tachyzoite produces an individual conductance transient

Transients were collected with the *T. gondii* wild-type (WT) RH strain using high bandwidth direct current measurements of host cells (COS1) under voltage-clamp conditions in the whole-cell configuration. Simultaneously, parasite invasion was monitored by DIC microscopy (Figure 1a). Multiple parasites were delivered to the host cells to increase the probability of capturing multiple invasions and the associated transient increases in whole-cell current. Invading tachyzoites showed a clear constriction (arrowheads, Figure 1a) and a single large current change was detected prior to the constriction of each parasite: multiple transients were recorded when multiple parasites invaded the same host cell (Figure 1b). Parasite invasion (*i.e.*, visible constriction) and the current transient were correlated events; no instance of invasion was observed without a transient (16 invasions with 16 transients). However, transients were also detected without seeing parasite invasion (9/25 WT transients), likely representing what has previously been described as abortive invasion (Koshy et al., 2012, Lamarque et al., 2014) and will be discussed later. Further analysis was performed on all the transients identified using the WT strain (n=25). The analysis revealed that every transient possessed characteristic wave form features – a fast rise to the peak and a slower fall to a new baseline – although the magnitude and duration of the transients varied (*e.g.,* Figure 1c and Supplementary Figure 1a and 1b). No obvious difference in transient conductance was seen based on the order of parasite entry, i.e. the first, middle, and last transients seemed similar (Figure 1c) suggesting that each transient represents an independent event irrespective of the number of invasions per cell. To corroborate independence, and because characteristic wave form features were similar, we evaluated all wild type-like conductance transient recordings, including those which were controls for our companion paper (see below and Male and Kegawa et al., 2024; Supplementary Figures 1a-h, 2a,b).

**Figure 1.**
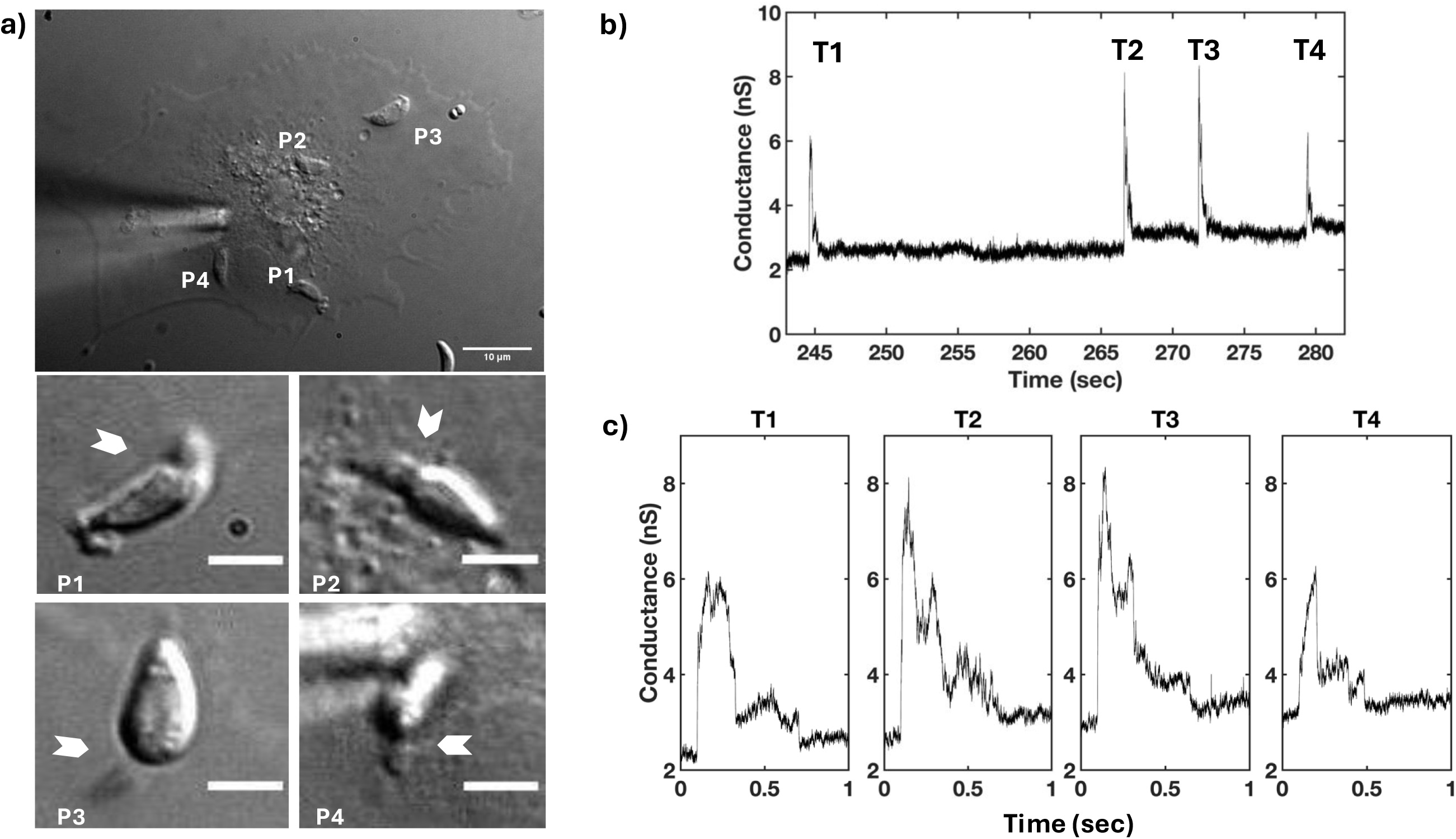
Each invading *T. gondii* WT RH strain tachyzoite produces a single electrical transient. a) DIC microscopy of *Toxoplasma* invasion of COS1 cell under visible patch pipette (upper panel); 10-µm scale bar. Four parasites (P1-P4) invaded during recording of a whole-cell patch-clamp electrophysiology experiment. White arrowheads indicate visible constrictions observed when parasites penetrate the host cell during invasion (middle panel: P1 and P2 invasions; lower panel: P3 and P4 invasions; 3-µm scale bar panels P1-P4). b) Multiple transients were detected from the electrophysiological recording obtained under the whole-cell configuration (-60 mV holding potential) during the invasions of the four parasites. c) Expanded time record of the transients (T1-T4) shown in b) for a clearer visualization of the wave form.

### The transient conductance increase induced by wild-type parasites occurs independently of complete moving junction formation

The depletion of rhoptry neck protein 2 (RON2) causes a severe early invasion defect in most of the parasites wherein rhoptry proteins are seen inside the host cell but the parasite subsequently detaches (Lamarque et al., 2014). However, a small number of the parasites can still invade through an alternative moving junction, meaning a moving junction without RON2 and its binding partners RON4 and RON5, but having some RON8 and perhaps other binding partners (Lamarque et al., 2014). To investigate whether RON2 insertion (and thus “complete” moving junction formation) is associated with the appearance of transients, we used both current recording experiments and the calcium influx assay (Male and Kegawa et al., 2024) to interrogate the ability of RON2 knockdown (KD-RON2) parasites to generate the conductance and calcium transients independently. KD-RON2 parasites do not express detectable levels of RON2, even without the ATc treatment typically used to induce knockdown, and show a severe invasion defect (Lamarque et al., 2014). While zero out of 159 KD-RON2 parasites invaded COS1 cells in our electrophysiology experiments, *i.e.,* no visible constrictions were observed by DIC microscopy, 19.5% (13.7%-26.5%; 95% CI) of the same parasites generated conductance transients (Figure 2a, Supplementary Figure 2a-b). Similar results were seen using the calcium transient assay (Figure 2a). KD-RON2 parasites generating conductance and calcium transients at levels similar to WT parasites stands in stark contrast to their near absence when the parasites are depleted of any of the individual proteins tested that regulate rhoptry exocytosis (Male and Kegawa et al., 2024). Together, the conductance and calcium transient results rule out the hypothesis that RON2 insertion into the host cell plasma membrane is itself the poration process. Furthermore, RON2-AMA1 binding (and by extension complete moving junction formation) are not required for the transient increase in host plasma membrane conductance.

**Figure 2.**
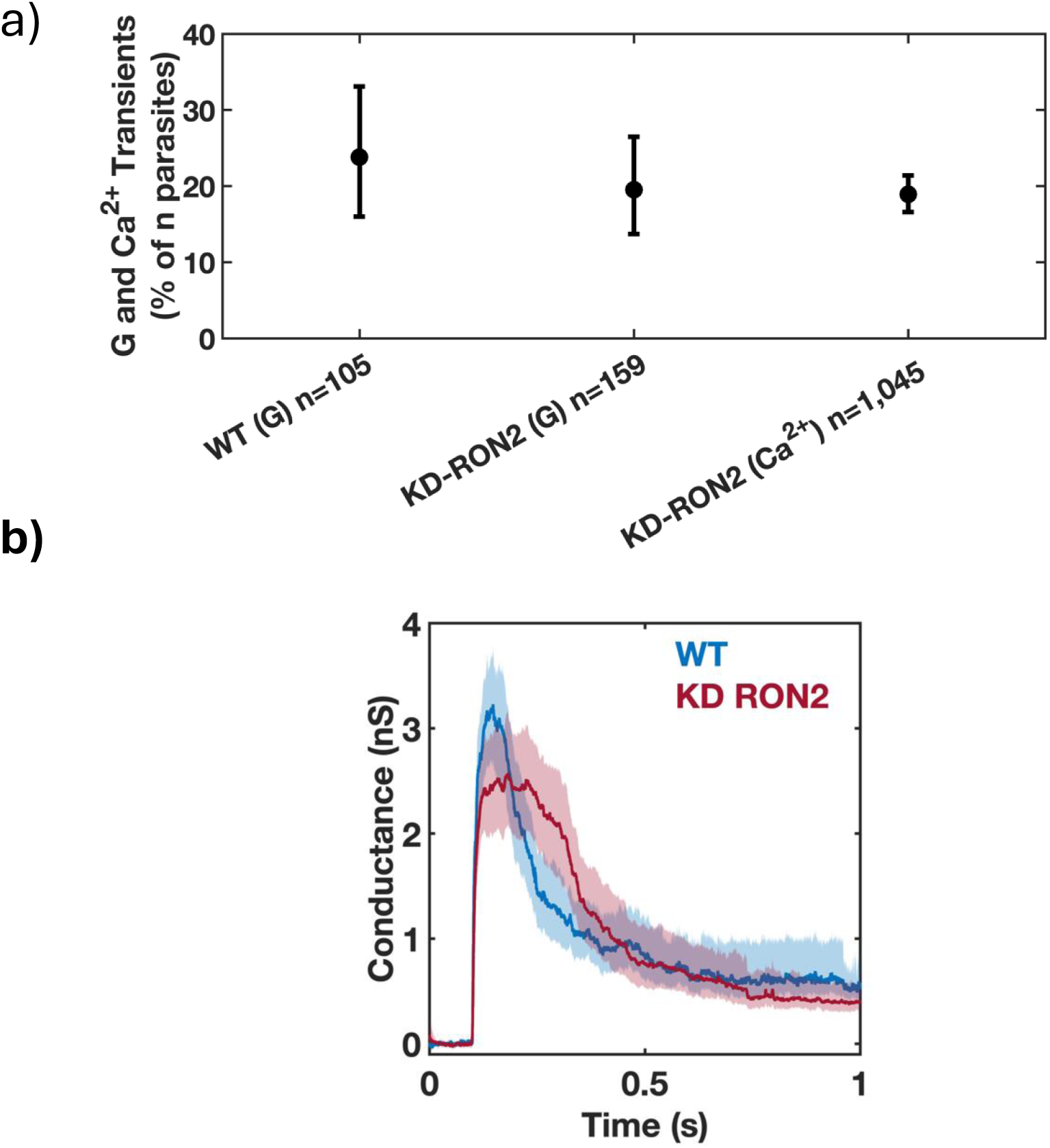
*T. gondii* invasion pore formation does not require RON2, and thus does not require complete moving junction formation. A) Phenotype of untreated WT and KD-RON2 parasites evaluated using electrophysiology (G) or calcium assay (Ca^2+^) data sets. Percentages of conductance and calcium transients (% of n parasites presented to host cells). Errors are Wilson, two-tailed upper and lower 95% confidence intervals. The percentage of transients observed in KD-RON2 (G) and KD- RON2 (Ca^2+^) are not significantly different from WT (G) (two-proportion z-test, z=0.8389, 1.227, p=0.40, 0.22, respectively). b) Conductance waveforms for WT (blue) and KD-RON2 strains (red) (mean, solid lines; 95% CI, shadings; n=25, 30, respectively).

### Detailed analysis of conductance transients induced by WT and KD-RON2 parasites

The conductance transients produced by both the WT (n=25) and untreated KD- RON2 parasites (n=30; hereafter in this paper referred to as KD-RON2) were averaged and compared (Figure 2b, Supplementary Figures 1a,b and 2a,b). On average, both WT and KD-RON2 parasite conductance transients show a rapid increase and then a slower decrease in conductance during the transient. To further characterize and compare the transients, change point analysis of individual transients was performed (see Methods and Figure 3). Individual transients are described by four characteristic features (mean±std. dev.) for WT and KD-RON2 (n=25 and n=30, respectively): 1) Peak conductance, calculated as the difference in conductance between the baseline and the maximum change point mean (WT: 3.40±1.12 nS and KD-RON2: 3.01±1.40 nS, Figure 4a); 2) Residual conductance, calculated as the difference in conductance between the pre- and post-transient baselines (WT: 0.54±0.38 nS and KD-RON2: 0.39±0.33 nS, Figure 4b); 3) Transient duration, calculated as the time between transient initiation and the beginning of residual conductance (WT: 322±219 ms and KD-RON2: 374±203 ms, Figure 4c), and 4) Peak conductance duration, calculated as the difference in the starting and ending changepoint time of the maximum conductance identified (WT: 73±64 ms and KD- RON2: 118±86 ms, Figure 4d). Both the WT and KD-RON2 transient waveforms begin with a rapid increase in conductance, reaching a maximum value with exponentially distributed time constants of 50±10 ms and 51±9 ms (mean±std. err.; n=25 and n=30), followed by exponentially distributed decreases (fall times) to the residual conductance with time constants of 197±39 ms and 203±36 ms (mean+/-std. err.; n=25 and n=30). For any one parasite strain (WT or KD-RON2), the rise time and fall time distributions are significantly different from each other; two-sample Kolmogorov-Smirnov test, *pWT*=3.97E-04 and *pKD-RON2*=8.08E-4; n=25 and n=30, respectively). Comparing the two strains, there is no evidence for differences between WT and KD-RON2 peak conductance, residual mean conductance, or mean transient duration. However, between WT and KD-RON2 strains the peak conductance duration is different (Figure 4d). The change point analysis supports the hypothesis that the kinetic process for creating and removing the conductance pathway for WT and KD-RON2 are similar (rise and fall time distributions, peak conductance, and transient duration: Figure 4a-c) but the lifetime (peak duration: Figure 4d) of the maximal conductance state is significantly longer in the absence of RON2.

**Figure 3.**
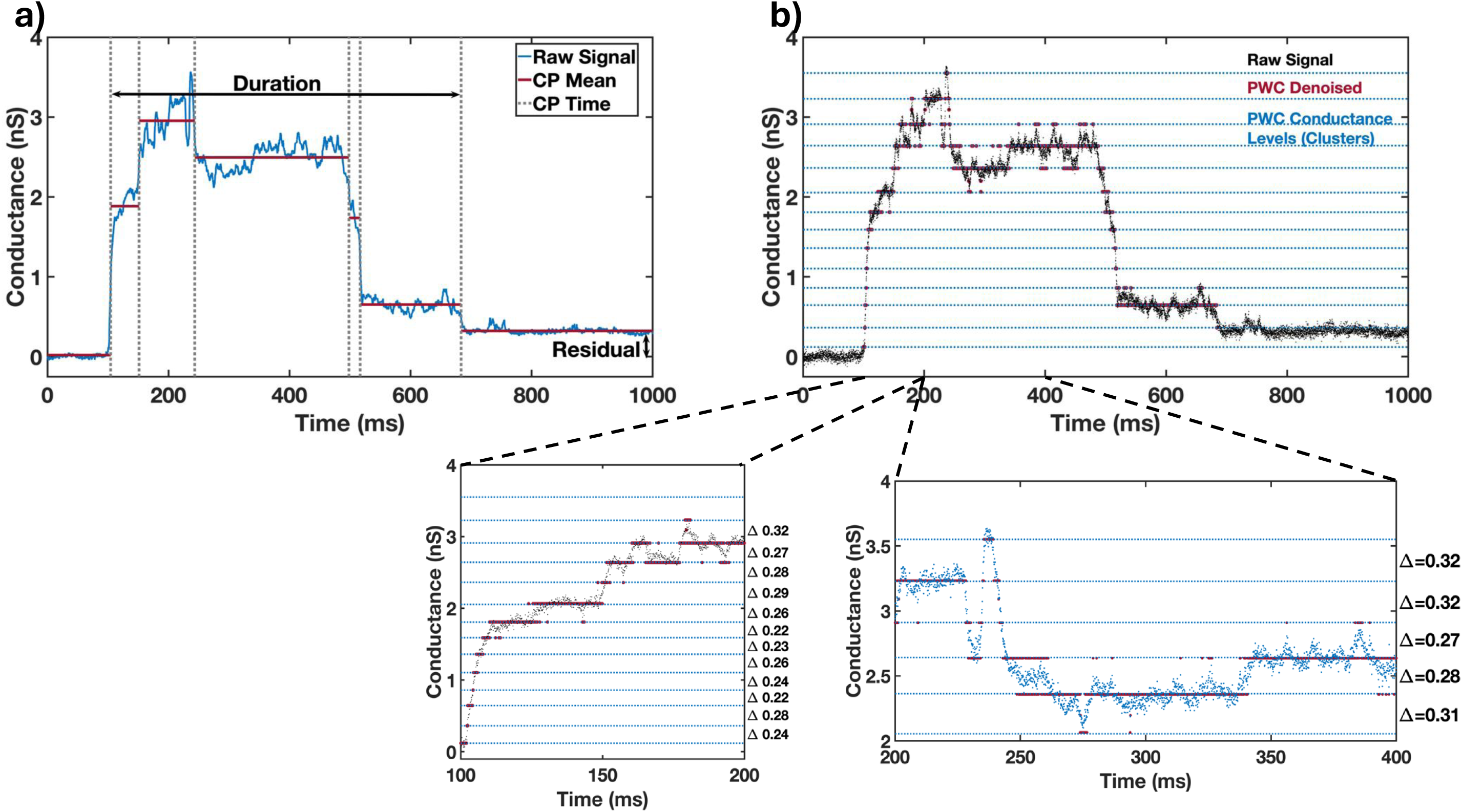
a) Representative Change Point Analysis (CPA) of a transient waveform induced by a WT parasite. Time dependent changes in conductance are represented by the gray dotted lines (CP Time) over a selected duration of 1 second. Characteristic features of the transient including the change point means (CP Mean, red), Duration, and Residual conductance (black arrows) are indicated. b) Representative PWC analysis of the same transient in a) showing the identified conductance levels (clusters) in blue. Expanded portions of the rising phase (100 ms) and a larger section of the transient (200 ms) are shown in the expanded plots. The changes in conductance between identified level sets (D) are indicated. The conductance changes from each set of transients were analyzed and used to identify the primary conductance change.

**Figure 4.**
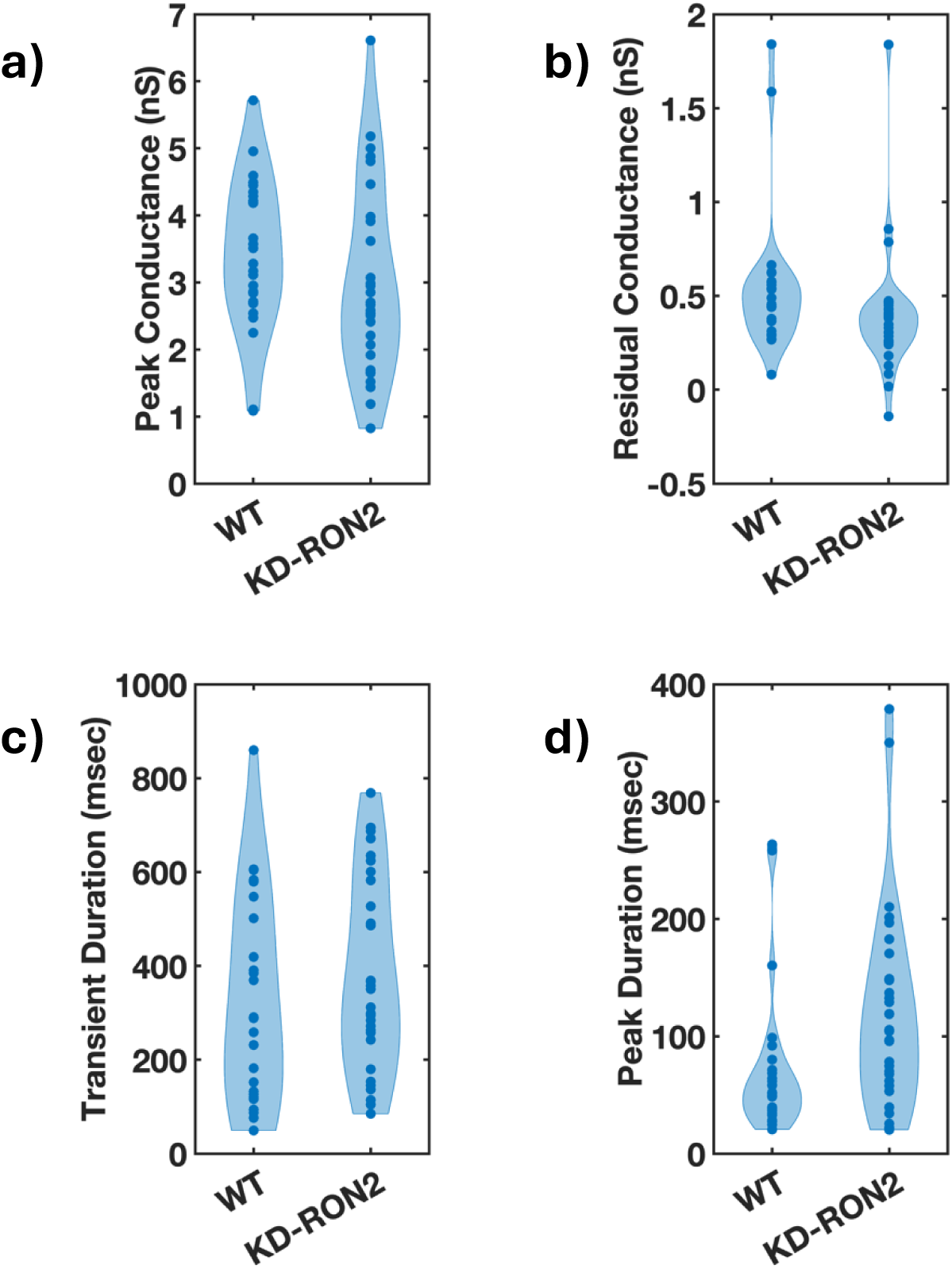
The conductance transients induced by WT and KD-RON2 tachyzoites differ in peak duration but not in transient duration nor peak or residual conductance. a-d). Violin plots comparing the waveform parameters obtained for WT and KD-RON2 parasites: a) peak conductance, b) residual conductance, c) transient duration, and d) peak conductance duration. Note, in 4d the peak durations are different; bootstrap differences in the mean, alpha=0.05; ANOVA, *p*=0.034 without Box-Cox transformation; following Box-Cox transformation and Welch’s T-test for unequal variance, *p*=5.69E-17.

### Step-like transients induced by WT and KD-RON2 parasites differ in quantal conductance

The above parameterization of the transients was consistent with similar pores produced by WT and KD-RON2 parasites, but the longer peak conductance duration observed in the averaged transients and confirmed in the analysis of the individual peak durations (Figures 2b and 4d) led to further investigation of the abrupt changes in conductance levels in both the rising (Supplementary Figures 1b, 2b) and falling phases (Supplementary Figure 1a, 2a) of the transients. To analyze the magnitudes and numbers of the rapid, step-like changes in conductance, each transient was processed using PWC denoising (Methods and Figures 3a, 5a,b). From the resulting conductance values the density distribution of the differences in adjacent conductance levels was obtained (Figures 3b, 5c). WT parasite-induced transients display a probability density peak at 0.26 nS, which can be considered a characteristic quantal step size (Figures 5a, 5c,d). This hypothesis was further tested by both parametric model fitting and Gaussian mixture modeling, each identifying a primary Gaussian peak at 0.26 nS with a width of 0.03 nS. The same analysis was applied to the KD-RON2 conductance transients, yielding a smaller primary Gaussian peak at 0.19 nS with a width of 0.03 nS (Figures 5b-d). The number of quantal units contributing to the averaged maximum conductance for both parasite lines are approximated by Poisson distributions with parameters 13.08 and 15.90 for WT and KD-RON2, respectively, an insignificant difference (no evidence using bootstrap differences in the Poisson parameter, alpha=0.05); Supplementary Figures 3a,b). Thus, knocking down RON2 does not eliminate the transient or the number of steps, but does change the step size significantly (27% smaller).

**Figure 5.**
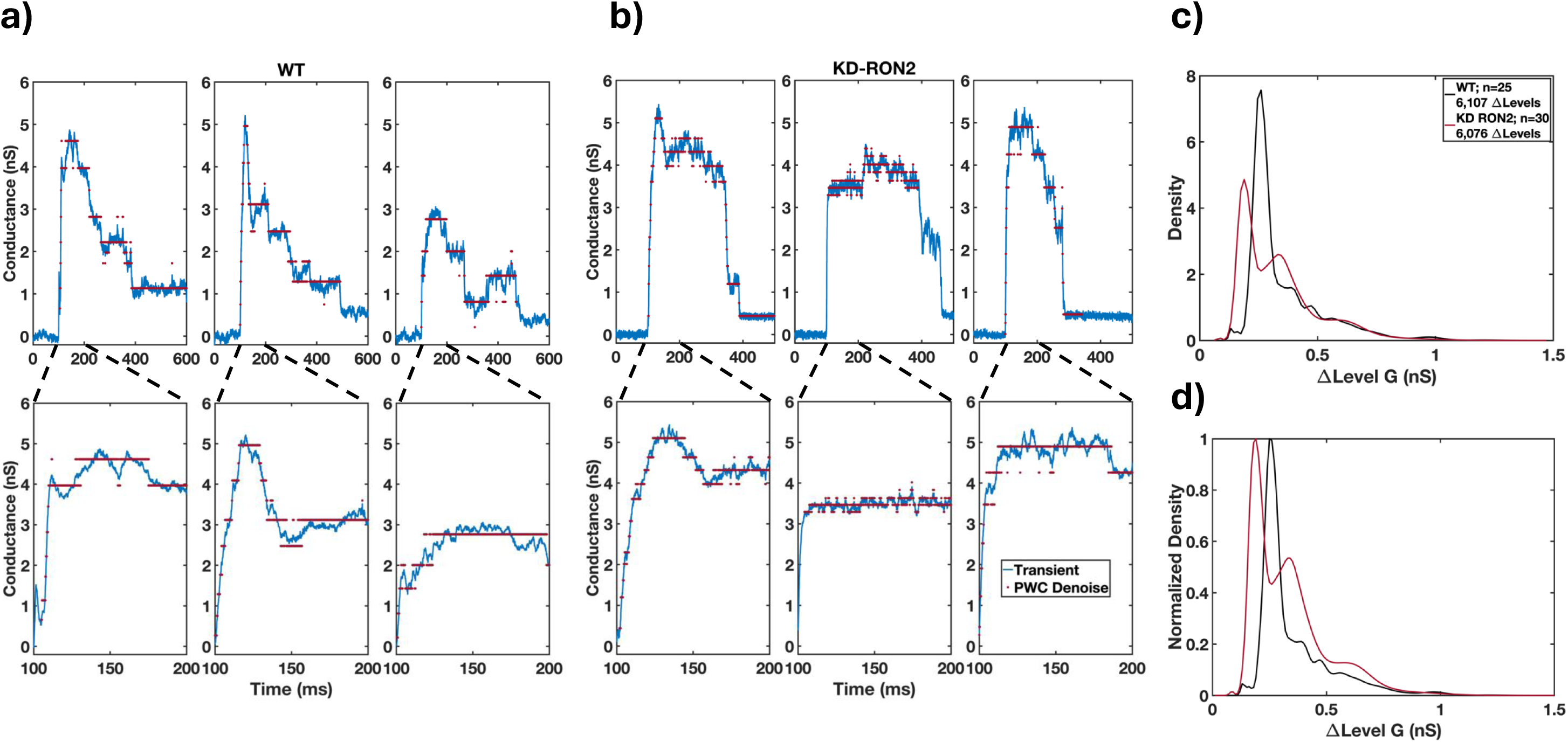
Evidence for quantal conductance levels from time series analysis of individual transient recordings. Piece wise constant (PWC) filtering was used to analyze the time-dependent conductance changes observed during individual transients. Examples of three a) WT-induced and b) KD-RON2 transients analyzed using PWC filtering to calculate the distributions of conductance level changes. c) Density distribution functions of combined transient conductance level changes (1Level G) produced by WT and KD-RON2 strains (n=25 and 30; respectively). d) Peak normalized density of data presented in c).

To control for the possiblility that the quantal step size parameter is overly sensitive, transient datasets generated by WT and other control parasites (WT parasites in low Ca^2+^ conditions and RASP2 conditional knockdown (cKD) parasites (Suarez et al., 2019) without anhydrotetracycline (ATc) treatment) were compared to increase our sample size. Ten additional transients were recorded in low Ca^2+^ (Supplementary Figures 1c,d), and 26 trasients using the cKD-TgRASP2 -ATc parasites (Supplementary Figures 2c,d). Both datasets showed the same characteristic wave form features as WT. PWC analysis was performed on each individual transient recorded and the density of conductance level changes were calculated. The primary peak quantal step values were 0.24 (0.03) nS and 0.25 nS (0.03) for WT low Ca^2+^ environment and cKD-TgRASP2 -ATc, respectively (mean (width)); the means are within the uncertainty of WT (0.26 (0.03)) consistent with conductance levels acquired from parasites having similar properties and similar quantal values. For completeness, all 4 groups were compared using both ANOVA (parametric) and Kruskal-Wallis (non-parametric) analyses. WT, WT low Ca^2+^, and cKD-TgRASP2 -ATc are significantly different from RON2 (Tukey’s p < 10^-15^, and Dunn’s post-hoc analyses, Q>21).

## DISCUSSION

The distinctive transient increase in host cell membrane conductance detected after parasite attachment but preceding the morphological changes of invasion implies that for a brief time ions can freely move across the host cell membrane. Most likely, this is due to the appearance in the host cell membrane of an aqueous pathway for ion flux, either due to protein-lined channels or a local breakdown of the lipid bilayer. The transient proceeds along a series of intermediates in conductance: once initiated, the conductance rises in tens of milliseconds, reaching a peak before diminishing more slowly to a residual conductance that is by one second after onset approximately 10% of the peak. Neither RON2 nor by extension the establishment of the RON2-AMA1 complex were required for the transient increase of either membrane conductance or Ca^2+^ entry. Also consistent with an aqueous pathway for ion flux (also known as a “pore”) is the proportionality of Ca^2+^ flux into the host cell cytoplasm during a transient with extracellular Ca^2+^ concentration, as shown in our companion paper, where it was also shown that transients require rhoptry exocytosis (Male and Kegawa et al., 2024). However, extracellular calcium is not itself likely to be the charge-carrying ion or required for either pore opening or closing, since conductance transients of similar shape and magnitude were observed in an environment with twenty times lower extracellular calcium concentration (Supplementary Figure 4). Based on these combined data, we propose that the conductance and calcium transients we have observed are manifestations of a rhoptry exocytosis-dependent host cell membrane poration process during invasion, whose purpose is to provide the pathway for rhoptry protein injection into the host cell cytosol. Consistent with this proposal are literature on abortive invasion: for KD-RON2, delivery of secreted rhoptry proteins to host cytosol persists without moving junction formation (Lamarque et al., 2014). What follows are deductions based on analysis of the electrophysiological data on this poration process and the development of the current working hypothesis (termed invasion pores, Figure 6).

**Figure 6.**
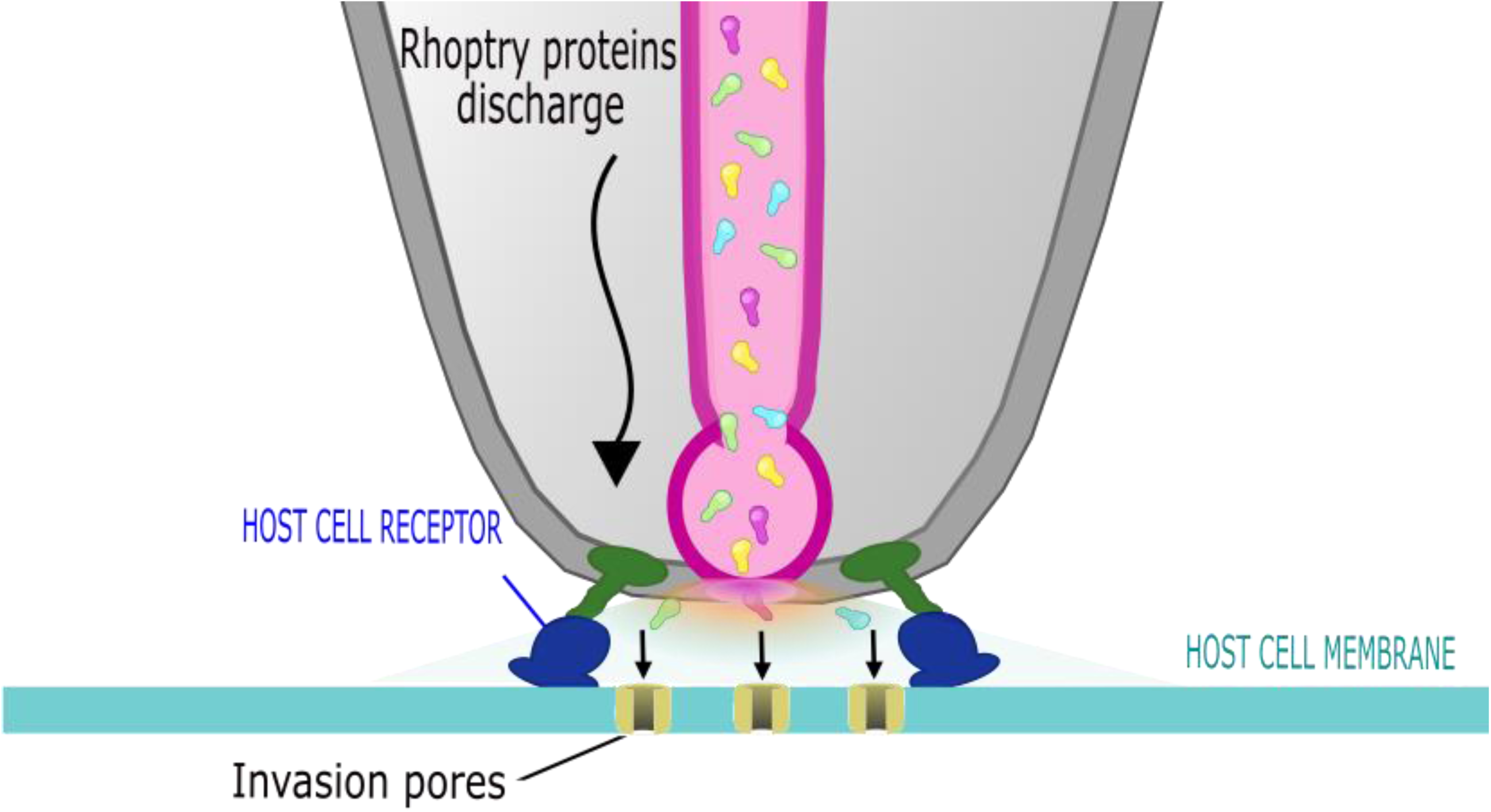
Model for invasion pore formation during parasite invasion. Invasion pore formation is triggered by rhoptry exocytosis (see companion paper, Male and Kegawa et al., 2024). Electrophysiological analysis supports multiple rather than single pore formation on the host cell membrane. Pore formation occurs prior to moving junction formation, but differences in the dominant quantal values induced by WT and KD- RON2 parasites suggest that RON2 contributes to the poration process or the passage of rhoptry contents through the pore.

### Kinetics of the conductance transient do not match one single large channel opening and closing

Every time invasion occurs, a transient conductance increase precedes the first invasion-associated morphological changes in the parasite. However, the role that the transient plays in invasion is unknown. Here we consider molecular structures that might underly the transient. The initial analysis of the data revealed that the average transient duration, peak conductance, and residual conductance were indistinguishable between WT and KD-RON2 parasites. This suggests that the underlying molecular structures for the transients are robust. The time dependence of the conductance change is not consistent with a simple model of one on/off channel opening and closing, which predicts one step increase in conductance followed by a step decrease of the same size. Not only are the recordings visually different from this prediction, but the transient relaxes to a residual conductance greater than the initial baseline, followed by a more variable time to return to baseline.

While instructive, comparing the WT transient maximum and quantal step conductances to ultrastructural size determinations is difficult. During *T. gondii* invasion, the only published size of a potential pore is 40 nm in diameter, measured from freeze-fracture electron microscopy images acquired after the moving junction has formed (Dubremetz, 2007). In this image, the host cell plasma membrane patch that had been in contact with the apical tip of the parasite and is now the presumptive PVM is smooth and continuous, other than the one circular structure. If a pore of this size was present and stayed open during all of invasion, the pore would have been detected in the time-resolved admittance measurements of the entire invasion process (Suss-Toby et al., 1996) but was not. If a physical cylindrical pore 40 nm in diameter were to form, the membrane conductance would be 34 nS and not the range of 1 - 6 nS observed here. Using a simple model to convert the peak conductance of the transients observed in our study to the diameter of a single cylindrical pore yields diameters of only 3 - 10 nm (Supplementary Figure 5). The freeze-fracture image might, however, depict a structure that contains the fused residua of several invasion pores.

### Multiple pores open and close in the host cell membrane during the invasion of Toxoplasma

The characteristics of the transient obtained from the high-resolution analysis suggest that multiple poration events occur to increase the permeability of the host cell membrane after rhoptry exocytosis and independent of complete moving junction formation. Detailed analysis of transients revealed a fast increase in the conductance with a slower decay, each composed of multiple “steps in conductance” identified by change point analysis (Figure 3a). Multiple steps in conductance are not unusual for proteinaceous pores having sub-conductance states, such as the voltage-dependent anion channel from Neurospora or rat liver (Zimmerberg & Parsegian, 1986). The largest component in the distribution of conductance step sizes for VDAC is the main open-closed transition (also ∼ 0.25 nS in salt solutions close to those used here).

Importantly, in the current study, the largest conductance component is different between WT (0.26 nS) and KD-RON2 (0.19 nS) parasites. This difference could result from RON2 either being directly involved in the structure of the invasion pore or regulating its formation or function. The lack of significant differences in the peak and residual conductance magnitudes, the time to reach both the peak and the residual conductance, and the transient duration of individual spikes (other than the duration of the maximal conductance level) indicates pore formation *per se* likely occurs irrespective of any residual RON2 and is not interfered with by an alternative moving junction complex containing at least RON8 (Lamarque 2014). If this alternative moving junction had more loosely apposed parasite and host membranes, allowing more ion diffusion across it, we would expect to report larger quantal values, not lower, so that is ruled out. Conversely if a moving junction lacking RON2 is more occluding to ion diffusion, this could explain these results. Also, if RON2 insertion is the direct cause of poration or if RON2 regulates pore opening, then the rise time of the transient would likely be different between WT and KD-RON2. However, there is no support for significant differences between WT and KD-RON2 when comparing their rise time or fall time distributions.

An alternative explanation for the difference in the distribution of step size between WT and KD-RON2 parasites is that depletion of RON2 modifies which rhoptry proteins pass through the pore, how they pass, or their local concentration near the invasion pore. Any of these factors could have an indirect impact on how ions flow through the pore, and might also explain the longer residence time of the KD-RON2 pore, indicated by the maximal conductance duration (Figure 4d). This result is understandable if the function of the invasion pore is related to the passage of rhoptry cargo through the pore: longer times are needed to move cargo through a smaller pore, and once the cargo has passed through, the invasion pore is no longer needed. In addition to the lack of RON2 in the KD-RON2 parasites, RON4 and RON5 are poorly expressed and mislocalized, and where a moving junction is observed it is comprised of alternative proteins such as RON8 (Lamarque et al., 2014). As documented above, the rising and falling phases are kinetically and distributionally indistinguishable for both WT and KD-RON2 transients, suggesting a RON2-*independent* insertion and removal process coupled to a RON2-*dependent* process that influences the lifetime of the maximal conductance state (Figure 2b).

### Known apicomplexan pores and translocons

If the pores described here function in rhoptry protein translocation, they may share features with other known protein translocons. In this study, the conductance transient induced during invasion is best fit by quantal increments of ∼0.26 nS, strikingly like the single channel values obtained by direct measurement for the pore-forming protein EXP2 (Garten et al., 2018), a critical component of the well-studied PTEX translocon in malaria parasites (de Koning-Ward et al., 2009, Kalanon et al., 2016). PTEX functions to export proteins across the parasitophorous vacuolar membrane (PVM) and into host red blood cell cytoplasm. The membrane conductance of EXP2 was measured using an on-vacuole patch clamp method, yielding a functional channel conductance of ∼0.2 - 0.3 nS (Garten et al., 2018).

In *T. gondii*, few pore-forming proteins or potential translocons have been identified. The PVM-associated Myr protein complex functions after invasion in the translocation of secreted dense granule proteins across the PVM into the host cell cytosol (Franco et al., 2016, Cygan et al., 2020). Dense granule proteins GRA17, GRA23, GRA47 and GRA72 are pore-forming proteins that also appear to function post-invasion in nutrient transport across the PVM (Gold et al., 2015, Bitew et al., 2024), although a role for these proteins in protein translocation across the PVM cannot be ruled out. Intriguingly, *Plasmodium* EXP2 can rescue loss-of-function phenotypes in *Toxoplasma* lacking GRA17 (Gold et al., 2015). *Toxoplasma* also encodes two perforin-like proteins, PLP1 and PLP2 (reviewed in Carruthers, Ann Rev Microbiol 2024). PLP2 is only expressed in the sexual stages (Kafsack et al., 2009) but PLP1 forms pores in the PVM and host cell membrane during tachyzoite egress from the host cell (Roiko & Carruthers, 2013). PLP1 pore-forming activity is pH-dependent, with higher activity in acidic compared to neutral conditions (Roiko & Carruthers, 2013). Invasion assays conducted in low pH conditions result in increased levels of parasite attachment and invasion, due to increased microneme secretion, and also increased levels of host cell wounding, potentially due to PLP1 (Roiko & Carruthers, 2013). Based on the structure of PLP1 (Ni et al., 2018), it seems unlikely it can function as a protein translocon.

The mechanism underlying the rapid closure of the invasion pore reported here is also a mysterious phenomenon. While not directly related to parasite invasion, membrane resealing in model systems of membrane repair is well documented, occurring on time scales from ∼1 to 10’s of seconds. (Steinhardt et al., 1994; Togo et al., 1999; Terasaki et al., 1997; Bansal et al., 2003; Humphrey et al., 2013; Klenow et al., 2021). Closing of the invasion pores may reflect 1) a host cell response to repair the sudden change in permeability through rapid endocytosis (Corrotte et al., 2020), 2) blebbing of the plasma membrane to remove the pores following their insertion (Jimenez et al., 2014), or 3) gated closure of the invasion pore once cargo is delivered. Finally, the existence of residual conductance implies that the invasion pore is not completely closed and remains in the host cell membrane throughout the invasion process, which takes a total of 15-20 sec.

## Conclusions

In this study we describe a transient permeability pathway in the host cell membrane during *Toxoplasma* invasion. The electrophysiology data are best explained by synchronous creation of multiple pores by material secreted from the rhoptries into the host cell plasma membrane (Figure 6). The function of these pores is unknown but their similar size to protein translocons and their appearance in the sequence of invasion (after rhoptry exocytosis and before moving junction formation) suggest that this poration creates a pathway for delivery of secreted rhoptry proteins into the host cell cytoplasm.

## MATERIAL AND METHODS

### Culture of fibroblast-like COS1 cells

Fibroblast-like COS1 cells (ATCC CRL-1650) were cultured in complete Dulbecco’s modified Eagle’s medium (DMEM supplemented with high glucose, 200 mM Glutamax, 1 mM sodium pyruvate, ThermoFisher Scientific, Cat#10569, Waltham, MA) with 10% Fetal Bovine Serum Premium Select (R&D Systems, Cat#S11550, Minneapolis, MN) and primocin 100 μg/mL (InvivoGen, Cat# ant-pm-2, San Diego, CA) at 37°C under 5% CO2. Cells were rinsed twice with DPBS without both CaCl2 and MgCl2 (ThermoFisher Scientific, Cat#14190, Waltham, MA). Cells were released from the flask using a 5-minute incubation with 0.25% trypsin containing 0.913 mM EDTA (GIBCO, Cat#25200-056, Waltham, MA). Trypsinized cells were centrifuged at 193 *x g*, 5 min, room temperature (RT), in complete DMEM, using an Allegra X-22R centrifuge with Beckman Coulter rotor. Cells were resuspended in complete medium and 1 mL containing 40,000 cells was seeded onto a 35 mm DT dish (Bioptechs, Butler, PA) for 60 minutes at 37°C under 5% CO2. Before starting electrophysiology experiments, DMEM medium was replaced by live cell imaging solution (LCIS) containing 155 mM NaCl, 3 mM KCl, 2 mM CaCl2, 1 mM MgCl2, 3 mM NaH2PO4, 10 mM HEPES, and 20 mM Glucose (final pH 7.4) or low-calcium LCIS containing 155 mM NaCl, 3 mM KCl, 0.1 mM CaCl2, 2.9 mM MgCl2, 3 mM NaH2PO4, 10 mM HEPES, and 20 mM Glucose (final pH 7.4).

### Culture of human foreskin fibroblasts (HFFs)

HFFs were maintained in Dulbecco’s Modified Eagle Medium (DMEM supplemented with high glucose, 200 mM Glutamax, 1 mM sodium pyruvate, ThermoFisher Scientific, Cat#10569, Waltham, MA) with 10% v/v heat-inactivated fetal bovine serum (FBS) (R&D Systems, Cat#S11550), 50 U/mL Penicillin-Streptomycin (Thermo Fisher Scientific, Cat#15140122, Waltham, MA) and 25 μg/mL gentamicin (GIBCO, Cat#15750, Waltham, MA). Prior to parasite passage, HFF medium was replaced with fresh culture medium (DMEM containing 10% v/v heat-inactivated FBS, 50 U/mL Penicillin-Streptomycin and 25 μg/mL gentamicin.

#### *Toxoplasma gondii* culture and isolation

*T. gondii* RH (hereafter designated WT) and KD-RON2 (Lamarque et al., 2014) strains were passaged in human foreskin fibroblasts (HFFs) (ATCC CRL-1634) grown in DMEM supplemented with 10% heat inactivated FBS (R&D Systems, Cat#S11550, Minneapolis, MN), 50 U/mL Penicillin-Streptomycin (Thermo Fisher Scientific, Cat#15140122, Waltham, MA), and 25 μg/mL gentamicin (GIBCO, Cat#15750, Waltham, MA) at 37°C under 5% CO2. *Toxoplasma* tachyzoites were isolated from infected monolayers at the large vacuole stage. Freshly released tachyzoites were removed and discarded before collecting infected cells from the culture flask by replacing culture medium with 4 mL of Endo buffer containing 89.4 mM KOH, 44.7 mM H2SO4, 10 mM MgSO4, 106 mM sucrose, 5 mM glucose, 20 mM Tris-H2SO4, and 3.5 mg/mL BSA (final pH 8.20, adjusted using KOH) (Endo et al., 1987). Attached cells and parasites were collected by scraping and resuspended in Endo solution. Collected infected cells were passed three times through a 25G needle and filtered through a 5-μm cellulose acetate filter (Sartorius, Cat#S7594-FMOSK, Göttingen, Germany). Isolated parasites were washed three times with Endo solution using a 10 min centrifugation at 400 *x g* and resuspended in 50 µL Endo solution for electrophysiological experiments. Parasites were used within two hours following isolation and were protected from light exposure.

### Calcium transient assay

#### *T. gondii* culture and isolation

For calcium transient experiments, “untreated” KD-RON2 parasites were grown in medium containing 0.07% ethanol for 48 hours prior to experiments for consistency with all other untreated controls reported in (Male and Kegawa et al., 2024). Isolated parasites were resuspended in anti-SAG1 antibody (monoclonal antibody DG52, a generous gift from Dr. David Sibley, 1/20 dilution of a 0.2 mg/mL stock) conjugated to AlexaFluor 647 (Alexa Fluor™ 647 Antibody Labeling Kit; Molecular Probes, Eugene, OR) in Endo buffer for 30 minutes at RT, then centrifuged at 1000*×g* for 2 minutes, and resuspended in Endo buffer at 3×10^7^ parasites/mL.

#### Cell labeling

HFFs were seeded in 3 chambers of an ibidi µ-Slide VI 0.4 (ibidi GmbH, Gräfelfing, Germany) for 2 hours at 37°C (with 5% CO2 and humidity) before indicator loading with 5 µM Fluo-4 AM (Invitrogen, Waltham, MA) with 1% PowerLoad Concentrate, 100X (Invitrogen, Waltham, MA) in LCIS for 90 minutes at RT.

#### Imaging and data analysis

Experiments were carried out at 35-36°C. Fluo-4 labeled cells were washed 1× with Endo buffer before allowing anti-SAG1 pre-labeled parasites to settle on the cells for 10 minutes. The buffer was then exchanged with pre-warmed (35-36°C) invasion-permissive LCIS to capture calcium transients and invasion events. Imaging was carried out on a Nikon Eclipse TE300 widefield epifluorescence microscope (Nikon Instruments, Melville, NY) using a 60× PlanApo λ objective (0.22 pixel/µm, NA 1.4). 1020×1020-pixel images were captured using an iXon 885 EMCCD camera (Andor Technology, Belfast, Ireland) set to trigger mode, with exposure time of 39 ms, no binning, 30 MHz readout frequency, 3.8× conversion gain, and 300 EM gain. Parasite induced perforation of host cells resulting in calcium transients and subsequent invasion events were observed by near-simultaneous excitation of Fluo-4 (490 nm) and AlexaFluor 647 (635 nm) using a pE-4000 LED illumination system (CoolLED, Andover England), through rapid excitation switching triggered by the NIS Elements Illumination Sequence module (Nikon Instruments, Melville, NY). Hardware was driven by NIS Elements v. 5.11 software (Nikon Instruments, Melville, NY). Calcium transient and invasion events were quantified as described elsewhere (Male and Kegawa et al., 2024) across three biological replicates, each consisting of three technical replicates for protein-depleted mutants and three technical replicates for controls, carried out on the same day. One technical replicate from one biological replicate was excluded from subsequent analysis due to poor quality.

### Electrophysiology

Electrophysiological experiments were performed with COS1 cells seeded onto a 35 mm DT dish containing LCIS, the external recording solution. Temperature during the experiment was kept at 37°C. The pipette solution for whole cell recordings contained 122 mM KCl, 2 mM MgCl2, 11 mM EGTA, 1 mM CaCl2, 5 mM HEPES (Final pH 7.26, adjusted with KOH). Patch pipettes (∼3 MΩ resistance) were fabricated from 1.5-mm thick wall borosilicate glass capillaries (P1000, Sutter Instruments, Novato, CA). Electrophysiological parameters of each patched cell were monitored with an amplifier (AxoPatch200b, Molecular Devices, San Jose, CA) in the voltage clamp mode. The output current was filtered using the internal 100 kHz Lowpass Bessel filter included in the amplifier, and an external 5 kHz low-pass 8-pole Bessel filter (Model 900 CT/9 L8L, Frequency Devices Inc, Haverhill, MA). A −60 mV holding potential was applied to monitor current changes due to interaction of each strain of *Toxoplasma* tachyzoites with COS1 cells. Current was recorded for a maximum of 15 minutes per cell, digitized at 100 μs (Axon Digidata 1550B and its associated software package Axopatch, Molecular Devices, San Jose, CA). Time resolution of <200 μs was empirically measured, matching the expected value determined by the interaction of the external 5 kHz filter and the whole-cell compensation circuitry of the amplifier. Conductance was calculated using Ohm’s law adjusted using the mean pipette access resistance, RA (n=40, RA=2.88/0.34 MΩ mean/sem); with our internal and external solutions, the measured reversal potential of the COS1 cells was ∼0 mV. Data analysis was performed offline (Clampfit 11.2, Molecular Devices, San Jose, CA and MatLab r2022b, MathWorks, Natick, MA).

### Tachyzoite delivery

Delivery pipettes were fabricated from 1.5-mm thick wall borosilicate glass capillaries using a pipette puller (P-80/PC, Sutter Instruments, Novato, CA). Delivery pipette inner diameters ranged from 10 to 20 µm, which was large enough to allow tachyzoites to pass through without damage. Delivery pipettes were bent (< 45°) over the flame of a small lighter to facilitate a vertical and smooth delivery of parasites onto the surface of a target cell. Parasites were delivered adjacent to the patched cell using a microinjector (FemtoJet® 4i, Eppendorf, Cat#5252000021D, Enfield, CT) set at ∼5 hPa.

### Optical visualization of invasion and image analysis

Imaging was initiated immediately after a stable, whole cell configuration was achieved. Parasite invasion was monitored on an inverted microscope (Axiovert 200, Zeiss, Oberkochen, Germany) equipped with a 63X 1.4 N.A. objective, differential interference contrast (DIC) optics, and a sCMOS camera (12.2 frames per second, 1501M-GE, ThorLab, Newton, NJ). Imaging proceeded for 15 min or until disruption of the giga-ohm seal required for whole cell electrophysiology, whichever occurred first. Offline, image files were screened for the presence or absence of parasite invasion, identified by DIC as a visible constriction of tachyzoites at the COS1 cell plasma membrane as they entered the cell. The number of parasites delivered onto the cell surface, as well as the number of invasions were counted to determine a ratio of invasion events per parasite.

### Data analysis for the electrophysiological recordings

Electrophysiological data was processed using MatLab. Digitized data were converted into a current and voltage data array using abfload (Hentschke, v1.4.0.0, MatLab Central File Exchange). Conductance and current plots were generated from the raw data following median filtering. MatLab median filter function, medfit1, was applied to remove noise spikes from the raw data to facilitate identification of biologically relevant transients. The choice of the median filter (9 points) was established empirically, balancing rejection of noise spikes, with minimal reduction in the amplitude of the parasite-induced transient. Typically, the noise spikes are significantly shorter than biologically relevant spikes with typical durations ∼ 0.4 ms. To identify the biologically relevant transients several criteria were used, including changes in the moving variance, first derivative, and manual inspection. The hand-curated selection of data was implemented in MatLab scripts to extract the transients. To identify transient and peak conductance starting and ending times, sequence change point analysis (MatLab findchangepts) was used to identify the times of mean conductance changes that persisted longer than 10 ms or 100 samples (100 μs each) (Figure 3a). Change point analysis identifies a change in signal statistics with time, for example the mean, of the whole cell conductance signal. As the number of change points existing in the conductance data was unknown, a range of fixed values were added to the residual error (a penalty term linear with the number of change points to compensate for the decrease in residual error that each additional change point introduced), until a plateau in the residual error was observed. The change point parameter set associated with stabilizing the residual error was used to calculate global features of the conductance transient: peak and residual conductance magnitudes, time to reach both the peak and residual conductance, and transient and peak duration. The peak is calculated from the magnitude of the maximum changepoint conductance value while the peak duration is the difference in time between the two change points defining the peak conductance. The transient duration is calculated from the difference between the first and last change points. The residual conductance is the difference between the mean of the last 50 ms and the mean of the first 95 ms in the 1 sec time series capturing each transient, where the first detected change in conductance for each spike is set at 100 ms. Time to reach the peak conductance is defined by the time between the start of the transient and the start of the maximum change point mean statistic and the time to reach the residual conductance is defined by the time between the end of the maximum change point mean and the last change point. Modeling the total resistance as the sum of the pore 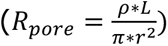 and twice the pore access 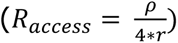 resistances, calculation of the radius (*r*) of a cylindrical pore (length *L*, solution resistivity *ρ*) with maximum conductance *G_max_* of each transient is

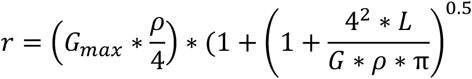

### Statistical analyses

Comparisons between the global features of the conductance transients observed in WT and KD-RON2 (peak and residual conductance, and transient duration) were evaluated using one-way analysis of variance (ANOVA) following both tests for normality (Lilliefors Test) and homogeneity of variance (Levene’s Test). The residual conductance distributions were not normal, displaying long-tails; one-way ANOVA of Box-Cox transformed data was used following the removal of one negative value in the KD-RON2 data set. To validate comparisons relying on distributional assumptions (one-way ANOVA), the bootstrap with 100,000 replicates was used to calculate 95% confidence intervals (CIs) at an alpha level of 0.05 for differences between data set means. CIs bracketing zero were consistent with no evidence for a significant difference between data sets. Rise and falling time distributions were compared using the two-sample Kolmogorov-Smirnov test. Rejection of the null hypothesis was set at p < 0.05 for ANOVA, Lilliefors, Levene’s, and Kolmogorov-Smirnov tests.

Two mixture model approaches were used to analyze distributions of data to provide a robust assessment: 1) fitting the empirical cumulative distribution function (eCDF) to a parametric model consisting of a weighted sum of normal or log-normal distributions and 2) applying a Gaussian mixture distribution model using both supervised (MatLab GMModel, iterative expectation maximization) and unsupervised (Wallace and Dowe, 2000 as coded by Statovic V 0.81, minimum message length) clustering algorithms. Graphs showing 95% confidence interval shadings were prepared using the Gramm data visualization toolbox for MatLab (V 2.27.1 by Pierre Morel). All statistical tests were two-sided unless indicated otherwise.

### Determination of clustered conductance values

Piece wise constant (PWC) filtering was used to analyze the time dependent conductance changes observed during a transient (Little & Jones, 2011; implemented in MatLab using pwc_cluster.m). PWC filtering is an unsupervised cluster analysis amenable to one-dimensional times series that optimizes membership within identified conductance levels (clusters). PWC filtering preserves sharp boundaries and other characteristics of rapid transitions (‘jumps’) in the data, and it does not require that the number of clusters be specified because the analysis is unsupervised. Here, a conductance level or state is hypothesized to be PWC with variable magnitude and duration depending upon the number and lifetime of individual conductance units present. Like change point analysis, removing noise from PWC signals is a signal processing optimization problem with many established iterative procedures. Here PWC filtering was applied using a clustering algorithm where each data point is shifted towards the highest density (mean-shift) within a fixed distance centered at the data point (hard kernel) (Little & Jones, 2011). In addition, the number of uniquely sized conductance levels following optimization is related to the amount of filtering and is set by an additional parameter called a ‘tuning hyperparameter’. Since the amount of filtering can be adjusted using this tuning hyperparameter, the PWC conductance levels are sensitive to the magnitude of this parameter, and there is a risk of fitting the noise if the parameter is too small. To avoid fitting the noise, an optimization procedure that identified the noise level of the baseline was adopted. The ∼100 ms baseline prior to the start of a transient (as above, it is set by us at 100 ms) was assumed to represent a PWC signal with noise properties associated with the whole cell configuration and that the noise present in the baseline is present and constant for the duration of the transient. The minimum constant conductance level change identified using the PWC filtering was set to be greater than the 99% confidence interval of the noise present in the constant baseline. The tuning hyperparameter was iteratively adjusted over a range until the resulting level changes were greater than the minimum constant conductance level change identified in the baseline noise. The hyperparameter satisfying the constraint defined by the baseline noise was considered optimal filtering for the transient. Since the noise varied between experiments, the resulting level changes were pooled from individually processed conductance transients.

### Testing for quantal conductance changes using the differences between adjacent clustered values

The magnitude of the level changes, obtained by calculating the absolute value of the differences between identified PWC levels present in the conductance transient time series, constitute the data sets whose distributional properties were analyzed as described above. To create the set of level changes, the PWC representation of the conductance time series was numerically differentiated by taking first order differences between the time series (ti-t(i-1)). Zero differences were dropped; only the positive and negative changes between conductance levels were analyzed. These changes represent the ‘jumps’ between levels at the specified time where the filtering defined a change in the cluster set associated with the new level. Except for sign, the positive and negative level changes had symmetric and equal distributions, justifying working with the absolute values of the level changes. If the distribution of level changes was uniform across all possible conductance differences (all possible conductance steps are represented in the data set), then there would be no evidence for a dominant peak or mode. However, if the distribution has one or more well defined peaks, then these peaks can be analyzed as a weighted sum of individual Gaussian distributions using mixture models to identify the peak with the greatest contribution to the overall distribution. Distributional peaks were visualized using a non-parametric density approximation prior to Gaussian mixture model analysis for identification and statistical evaluation of the peak properties, as described above. Comparisons between modes of multi-modal distributions were made by first isolating sub-distributions of data using the 95% CI around the Gaussian mixture model identified peaks followed by pair-wise and group analyses (ANOVA and Kruskal-Wallis) and subsequent post-hoc multiple comparisons testing using Tukey’s Honestly Significant Difference or Dunn’s test, respectively.

## ACKNOWLEDGEMENTS

Transgenic parasites were kindly gifted from Dr. Maryse Lebrun from Université Montpellier, whom we thank for useful discussions. This work was supported by the intramural program of the NICHD (JZ) and U.S. Public Health Service grant U01AI169067 (GEW). YK was supported in part by a JSPS research fellowship for Japanese Biomedical and Behavioral researchers at NIH from 2021 to 2023. The authors declare that they have no conflict of interest.

## AUTHOR CONTRIBUTIONS

Experiments were performed by YK, FM, IJM and EM, data analysis PB, YK and FM, conceptualization and writing was conducted by all authors.

## SUPPLEMENTARY FIGURE LEGENDS

**Supplementary Figure 1a.** Gallery of 25 WT parasite conductance transients calculated from recorded current measured using −60 mV holding potential in an external buffer containing 2.0 mM CaCl2. The initial 100 ms of baseline is plotted prior to the detection of the transient.

**Supplementary Figure 1b.** Gallery of 25 WT parasite conductance transients calculated from recorded current measured using −60 mV holding potential in an external buffer containing 2.0 mM CaCl2. The initial 5 ms of baseline prior to the detection of the transient and the initial 105 ms of each recorded transient are plotted.

**Supplementary Figure 1c.** Gallery of 10 WT parasite conductance transients in low external calcium, calculated from recorded current measured using −60 mV holding potential in an external buffer containing 0.1 mM CaCl2. The initial 100 ms of baseline is plotted prior to the detection of the transient.

**Supplementary Figure 1d.** Gallery of 10 WT parasite conductance transients in low external calcium, calculated from recorded current measured using −60 mV holding potential in an external buffer containing 0.1 mM CaCl2. The initial 5 ms of baseline prior to the detection of the transient and the initial 105 ms of each recorded transient are plotted.

**Supplementary Figure 1e.** Gallery of 26 cKD_TgRASP2 -ATc parasite conductance transients calculated from recorded current measured using −60 mV holding potential in an external buffer containing 2.0 mM CaCl2. The initial 100 ms of baseline is plotted prior to the detection of the transient. Because no ATc or solvent is applied, aside from the genetic alteration of the parasite no change of the protein complement of this parasite line compared to WT is expected.

**Supplementary Figure 1f.** Gallery of 26 cKD_TgRASP2 -ATc conductance transients calculated from recorded current measured using −60 mV holding potential showing the initial 105 ms of the transients. 5 ms of baseline is included prior to detection of the transient. Because no ATc or solvent is applied, aside from the genetic alteration of the parasite no change of the protein complement of this parasite line compared to WT is expected.

**Supplementary Figure 2a.** Gallery of 30 KD-RON2 parasite conductance transients calculated from recorded current measured using −60 mV holding potential in an external buffer containing 2.0 mM CaCl2. The initial 100 ms of baseline is plotted prior to the detection of the transient.

**Supplementary Figure 2b.** Gallery of 30 KD-RON2 parasite conductance transients calculated from recorded current measured using −60 mV holding potential showing the initial 105 ms of the transients in an external buffer containing 2.0 mM CaCl2. The initial 5 ms of baseline is plotted prior to the detection of the transient.

**Supplementary Figure 3.** The same number of quantal units contribute to the conductance maxima of WT and KD-RON2 transients. a) Transformation of the mean transient conductance (Figure 3a) using the peak quantal sizes for WT (0.26 nS) and KD-RON2 (0.19 nS), rounded to nearest integer. b) Violin plots of the number of quantal units at the maximum conductance of WT and KD-RON2 transients (n = 25 and 30, respectively).

**Supplementary Figure 4.** Transients induced by WT parasites in low external calcium concentration (0.1 mM CaCl2) have similar properties to WT transients induced in regular LCIS (2.0 mM CaCl2; Figure 3a). The average waveform of low-calcium transients displays the characteristic fast rise to peak conductance and slower recovery to a new baseline (n=10). The confidence interval (CI) is calculated for each point along the averaged transient.

**Supplementary Figure 5.** Violin plot of pore diameters calculated from individual WT transient maximum conductance using a model for a cylindrical pore (see methods).

## Notes

### Competing Interest Statement

The authors have declared no competing interest.

### Summary of Updates

The new version of the manuscript includes a larger dataset and more detailed analysis of the parasite strains used in the study for a better comparison of the discrete changes of the quantal values in the raise and closure phases of the transients described along the manuscript. This allows a better description of the moving junction during parasite invasion since the reduced value of the transients caused by KD-RON2, compared to the transients caused by wild type parasites, rule out a direct structural role of RON2 protein in the pore assembly or its role in restricting the diffusion of the pore-forming components.

