## Supplementary Figures for "The invasion pore induced by *Toxoplasma gondii*"

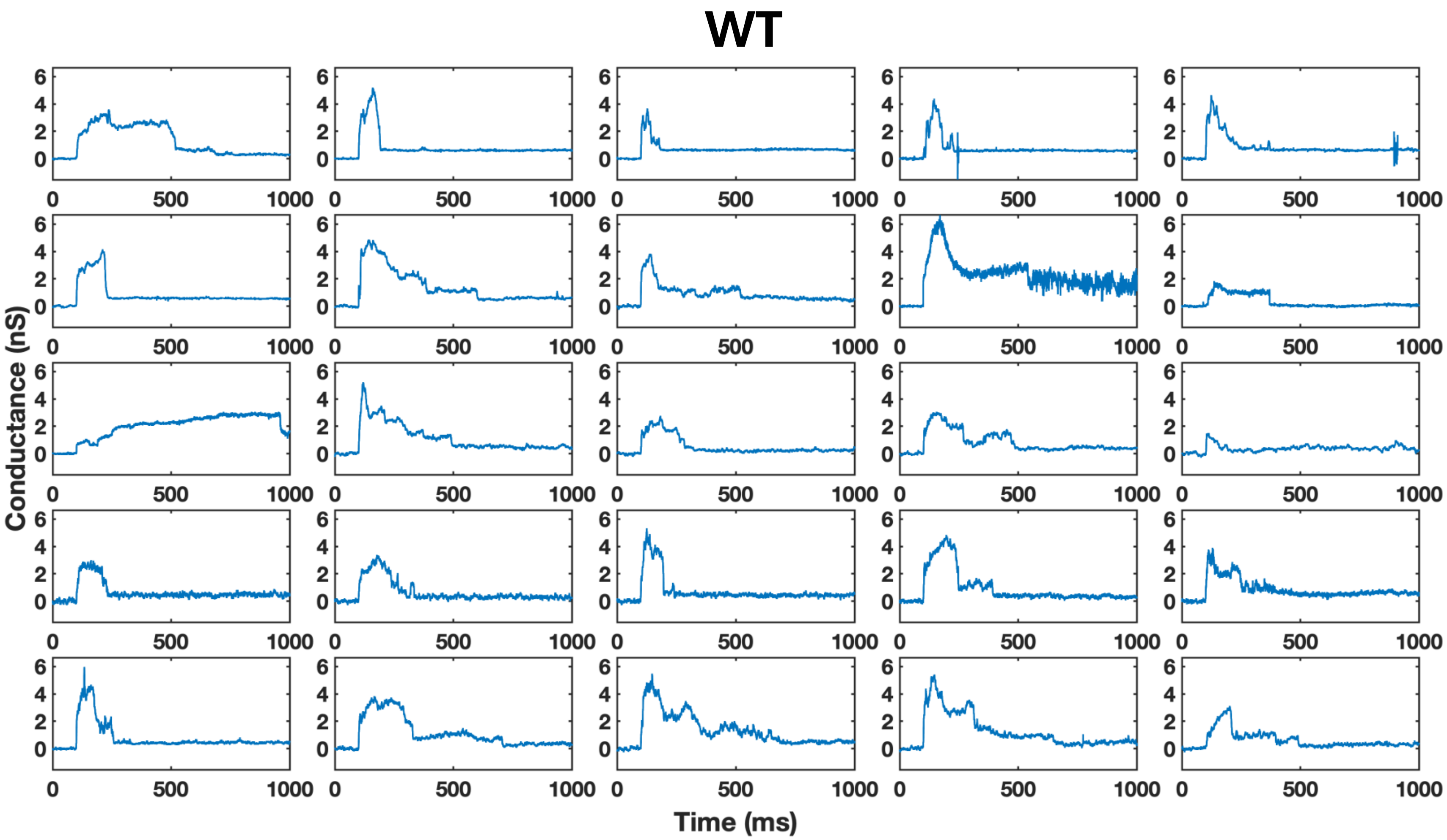


**Supplementary Figure 1a.** Gallery of 25 WT parasite conductance transients calculated from recorded current measured using -60 mV holding potential in an external buffer containing 2.0 mM $CaCl_2$. The initial 100 ms of baseline is plotted prior to the detection of the transient.

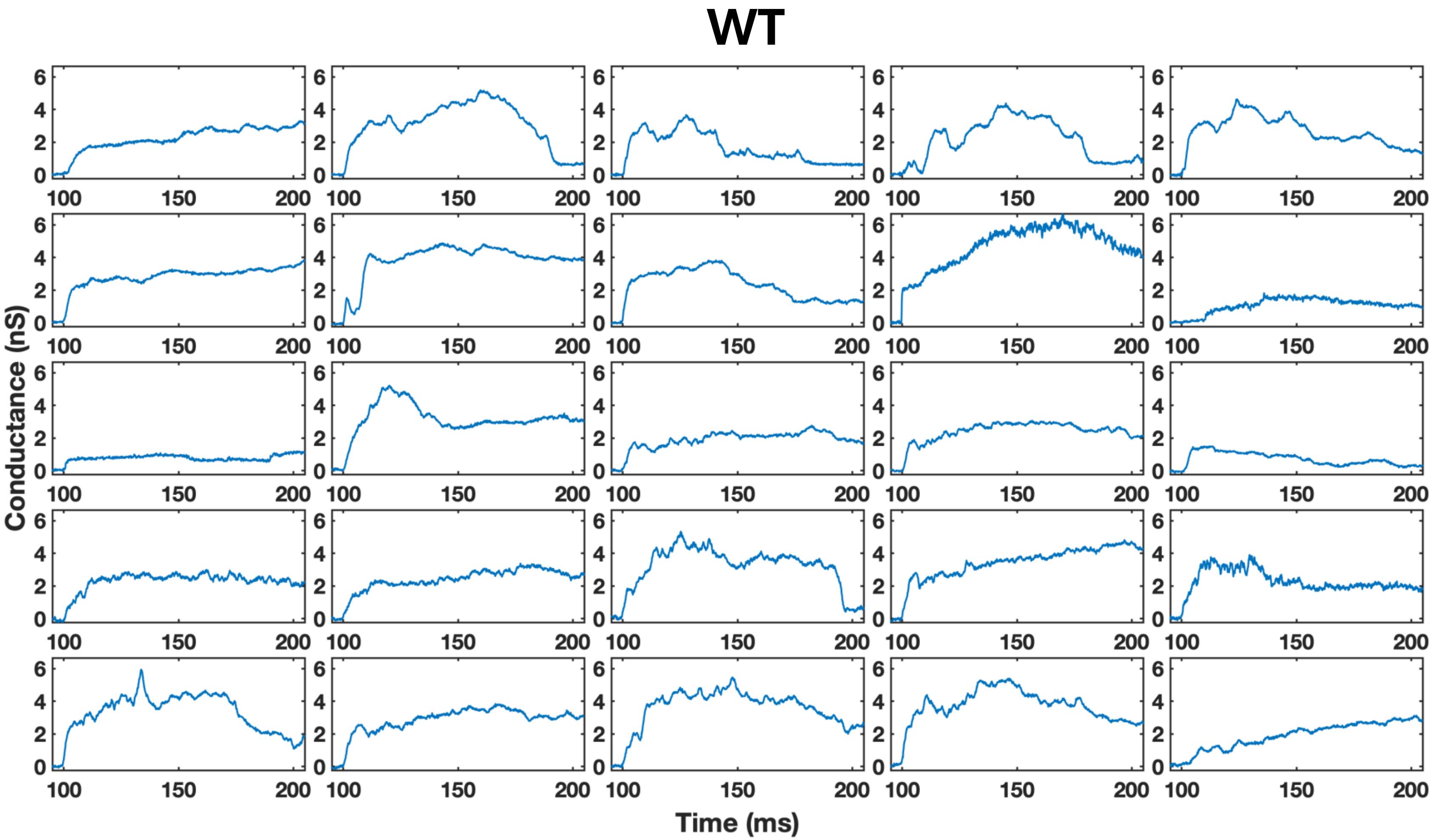


**Supplementary Figure 1b.** Gallery of 25 WT parasite conductance transients calculated from recorded current measured using -60 mV holding potential in an external buffer containing 2.0 mM $CaCl_2$. The initial 5 ms of baseline prior to the detection of the transient and the initial 105 ms of each recorded transient are plotted.

### Low $Ca^{2+}$ WT

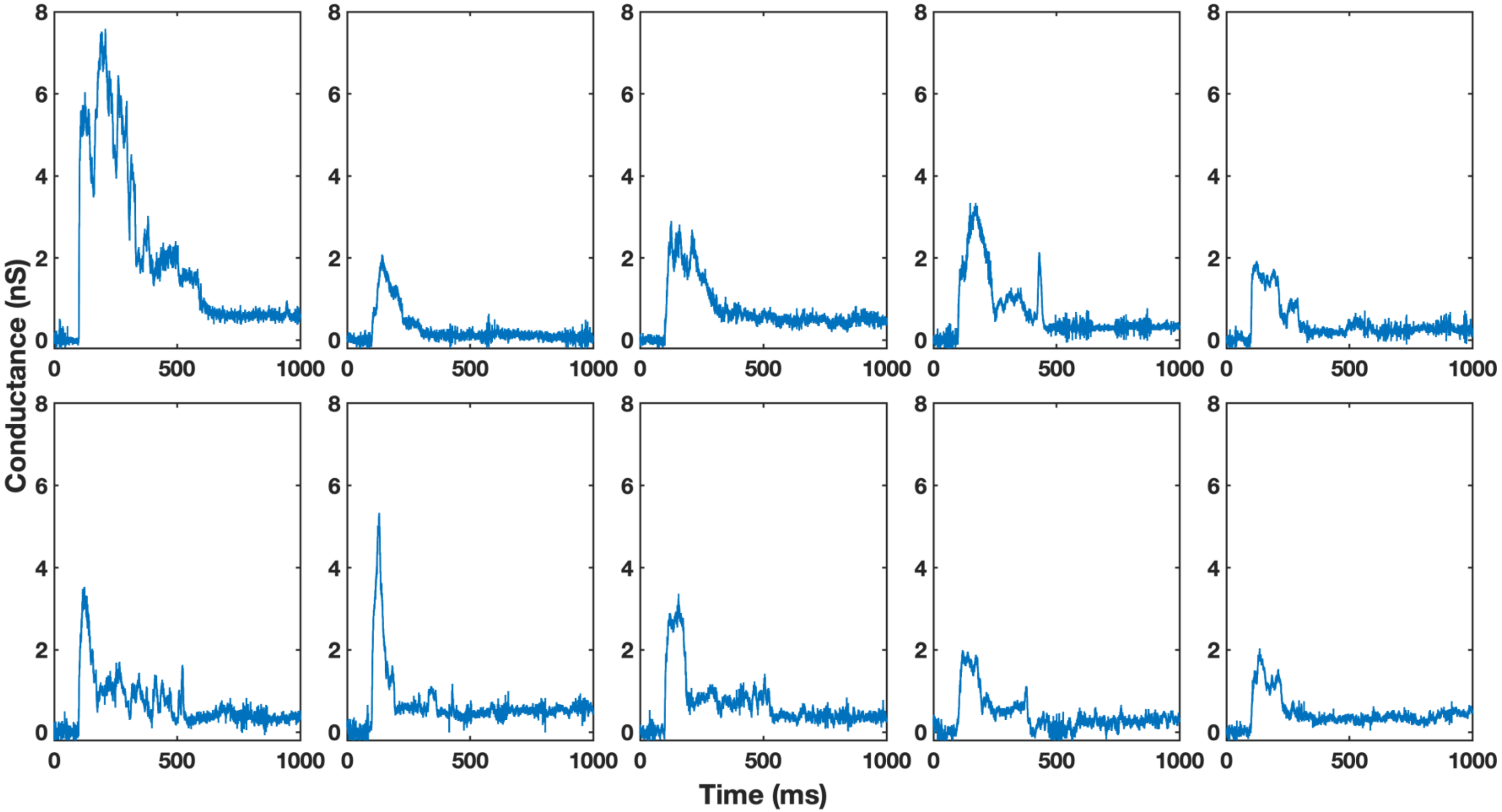


**Supplementary Figure 1c.** Gallery of 10 WT parasite conductance transients in low external calcium, calculated from recorded current measured using -60 mV holding potential in an external buffer containing 0.1 mM $CaCl_2$. The initial 100 ms of baseline is plotted prior to the detection of the transient.

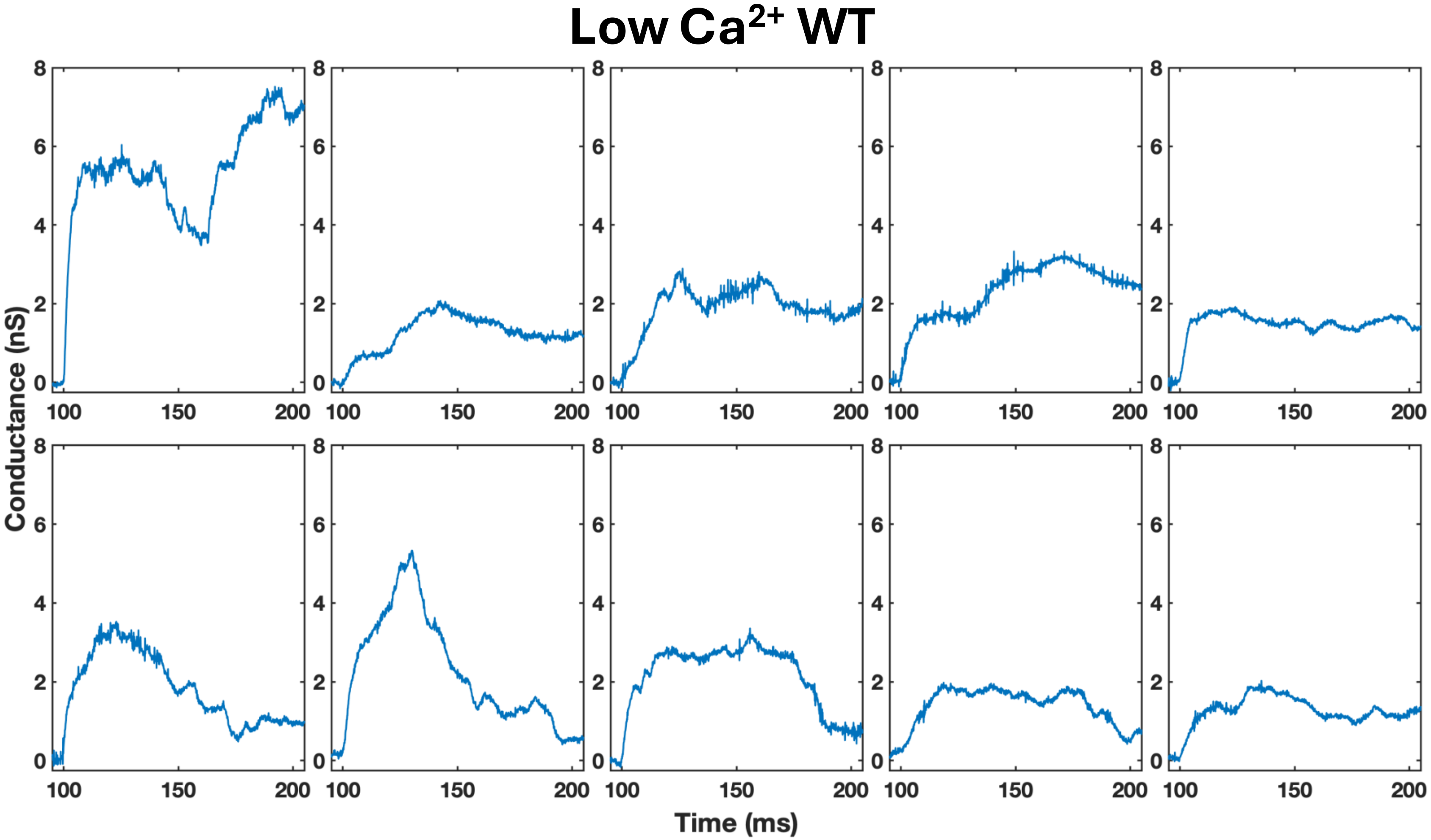


**Supplementary Figure 1d.** Gallery of 10 WT parasite conductance transients in low external calcium, calculated from recorded current measured using -60 mV holding potential in an external buffer containing 0.1 mM $CaCl_2$. The initial 5 ms of baseline prior to the detection of the transient and the initial 105 ms of each recorded transient are plotted.

### cKD_TgRASP2 -ATc

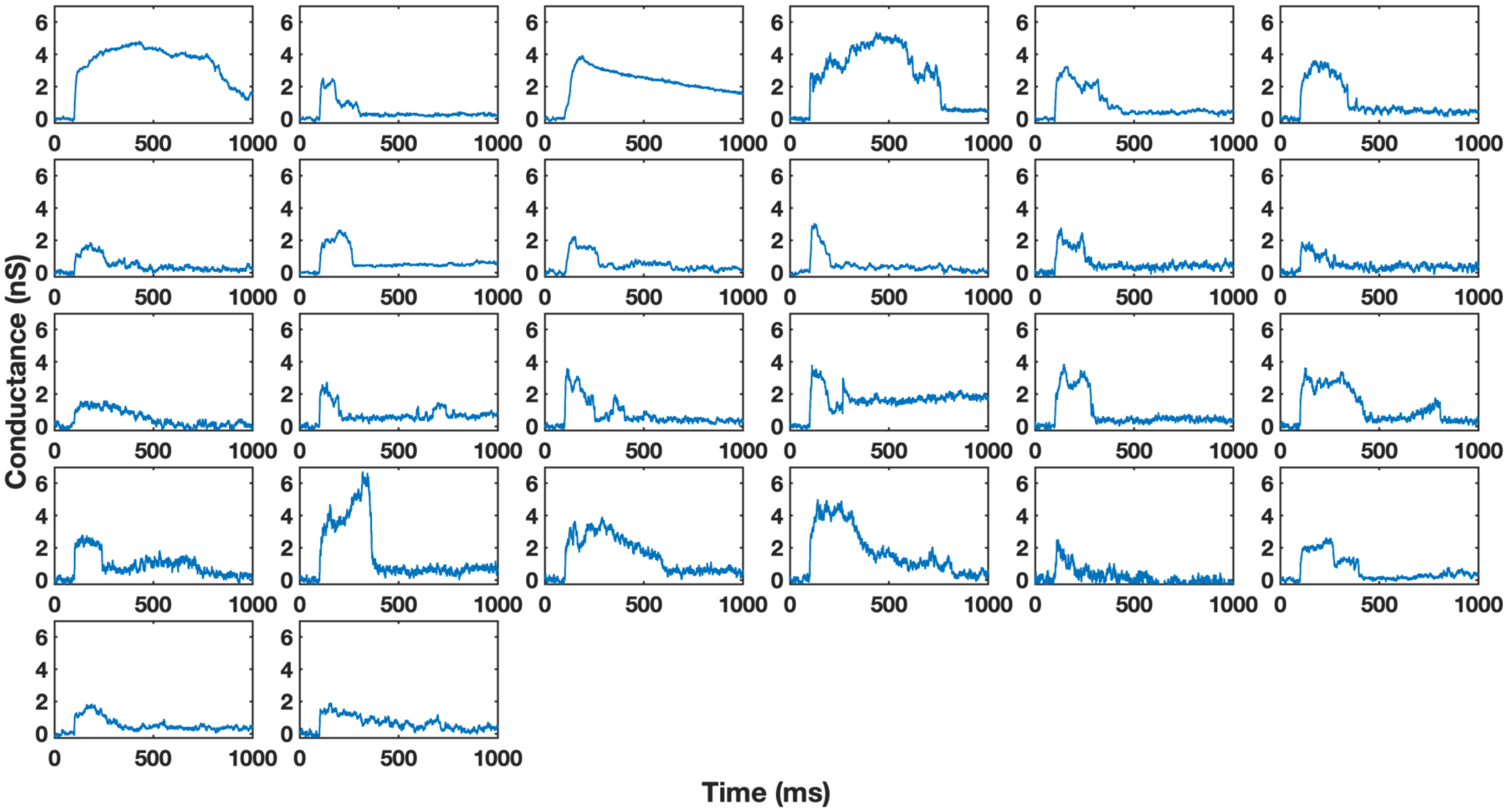


**Supplementary Figure 1e.** Gallery of 26 cKD_TgRASP2 -ATc parasite conductance transients calculated from recorded current measured using -60 mV holding potential in an external buffer containing 2.0 mM $CaCl_2$. The initial 100 ms of baseline is plotted prior to the detection of the transient. Because no ATc or solvent is applied, aside from the genetic alteration of the parasite no change of the protein complement of this parasite line compared to WT is expected.

### cKD_TgRASP2 -ATc

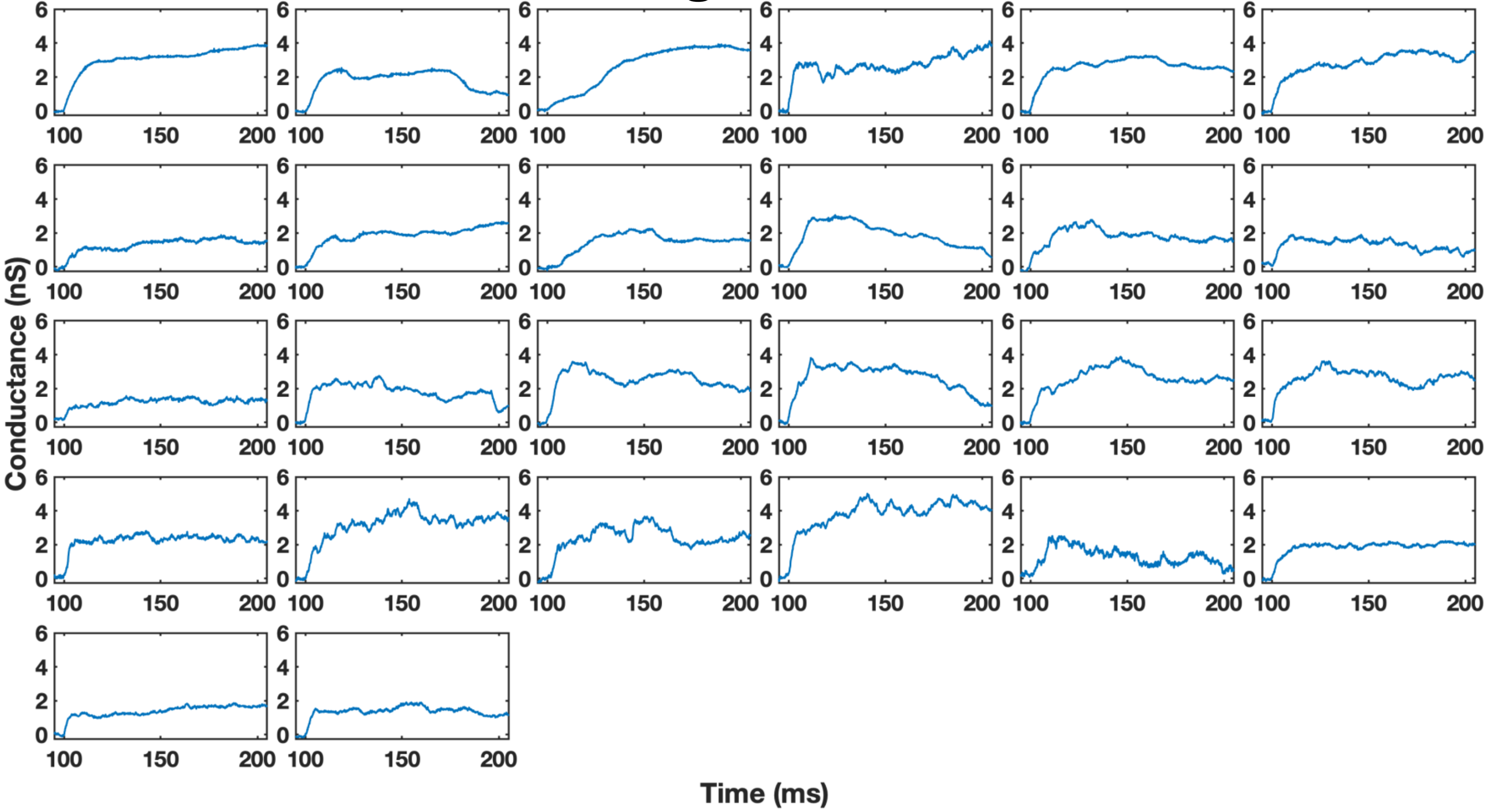


**Supplementary Figure 1f.** Gallery of 26 cKD_TgRASP2 -ATc conductance transients calculated from recorded current measured using -60 mV holding potential showing the initial 105 ms of the transients. 5 ms of baseline is included prior to detection of the transient. Because no ATc or solvent is applied, aside from the genetic alteration of the parasite no change of the protein complement of this parasite line compared to WT is expected.

### KD RON2

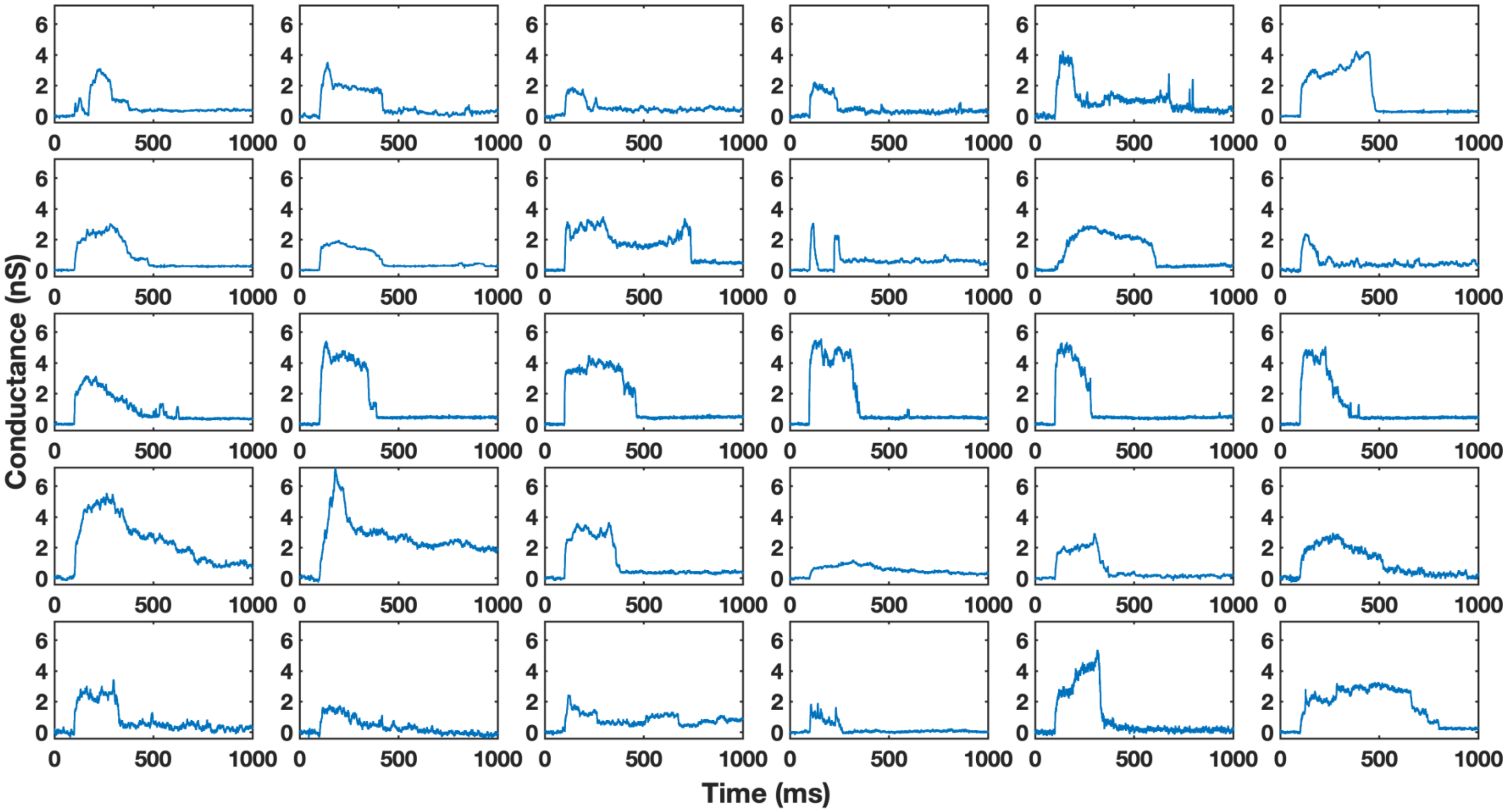


**Supplementary Figure 2a.** Gallery of 30 KD-RON2 parasite conductance transients calculated from recorded current measured using -60 mV holding potential in an external buffer containing 2.0 mM $CaCl_2$. The initial 100 ms of baseline is plotted prior to the detection of the transient..

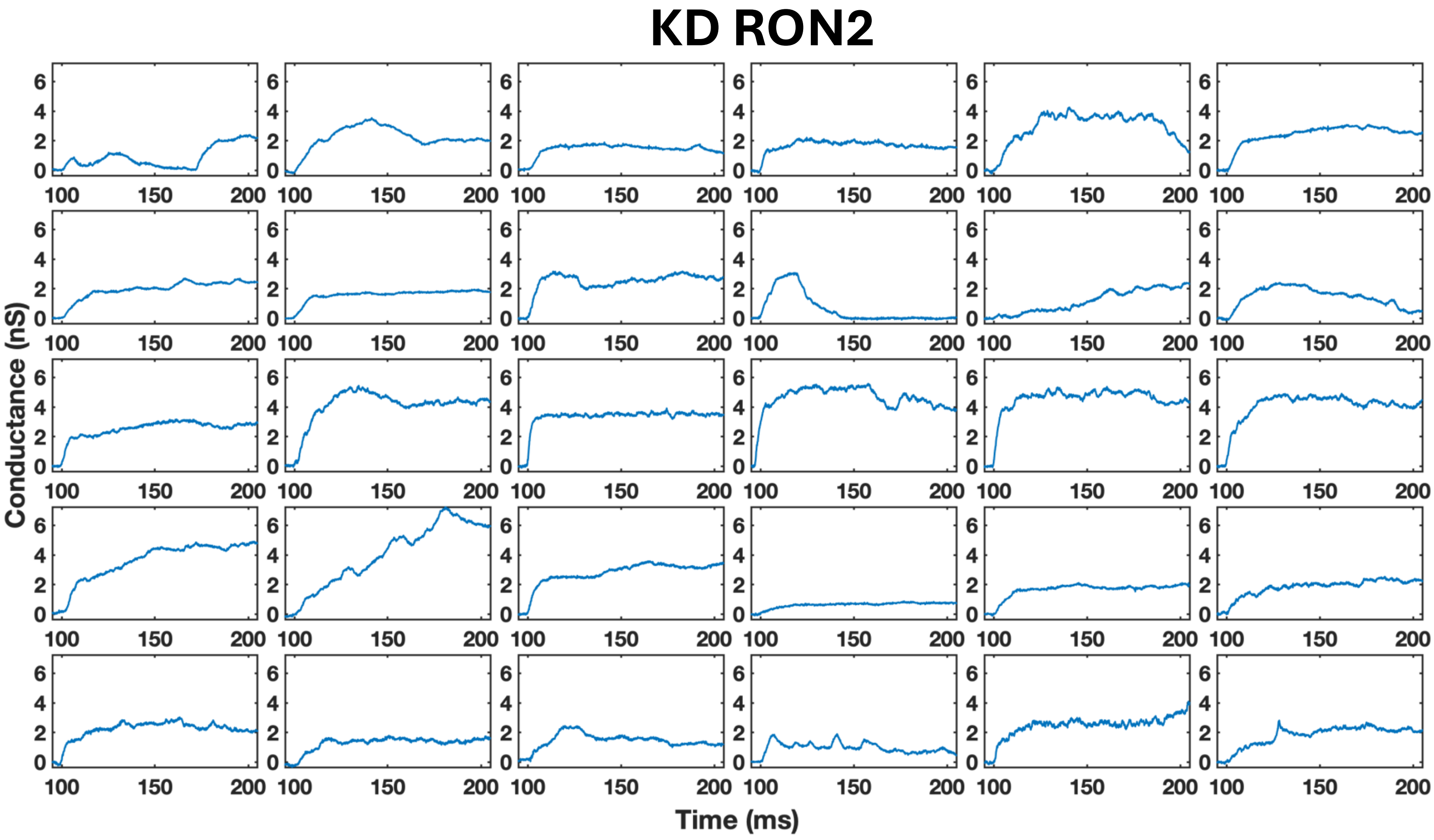


**Supplementary Figure 2b.** Gallery of 30 KD-RON2 parasite conductance transients calculated from recorded current measured using -60 mV holding potential showing the initial 105 ms of the transients in an external buffer containing 2.0 mM $CaCl_2$. The initial 5 ms of baseline is plotted prior to the detection of the transient.

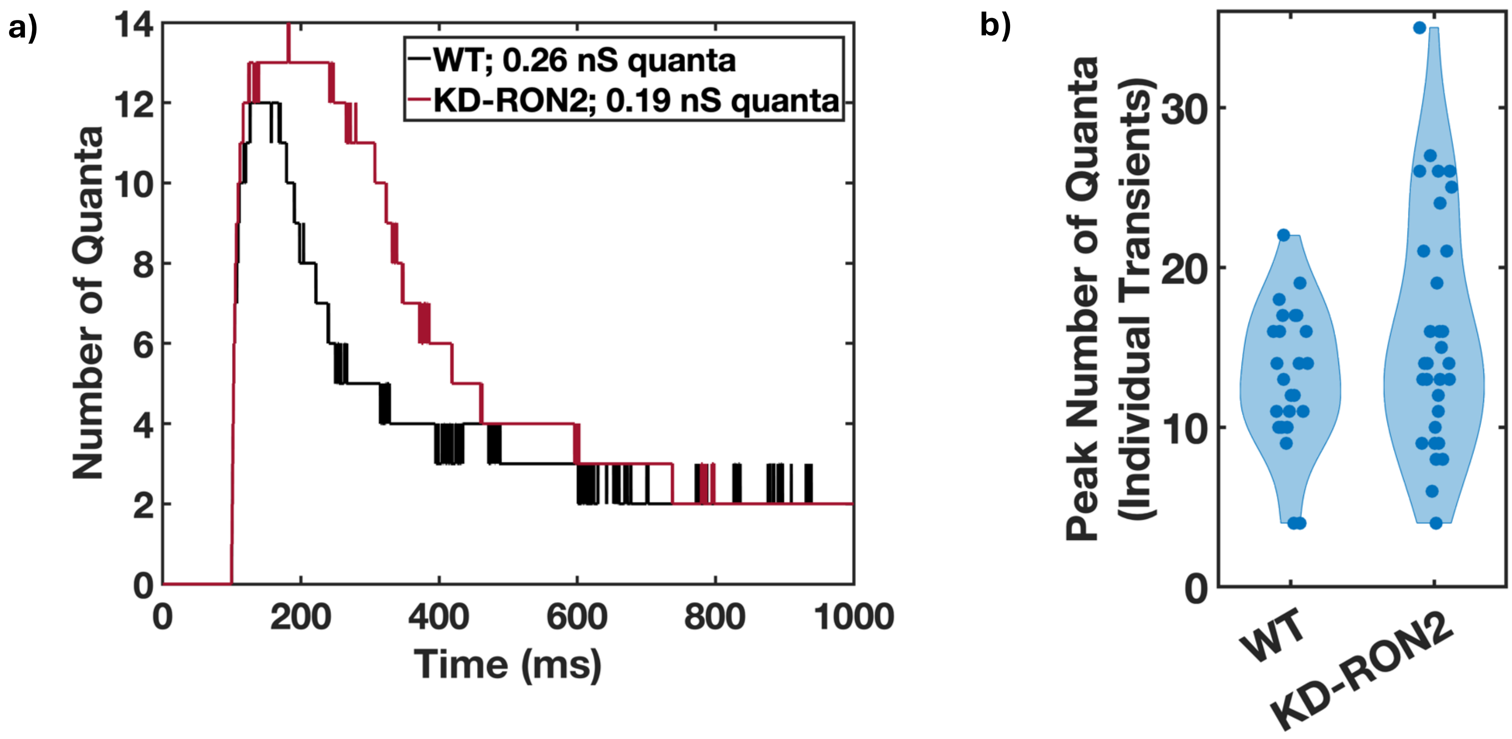


**Supplementary Figure 3.** The same number of quantal units contribute to the conductance maxima of WT and KD-RON2 transients. a) Transformation of the mean transient conductance (Figure 3a) using the peak quantal sizes for WT (0.26 nS) and KD-RON2 (0.19 nS), rounded to nearest integer. b) Violin plots of the number of quantal units at the maximum conductance of WT and KD-RON2 transients (n = 25 and 30, respectively).

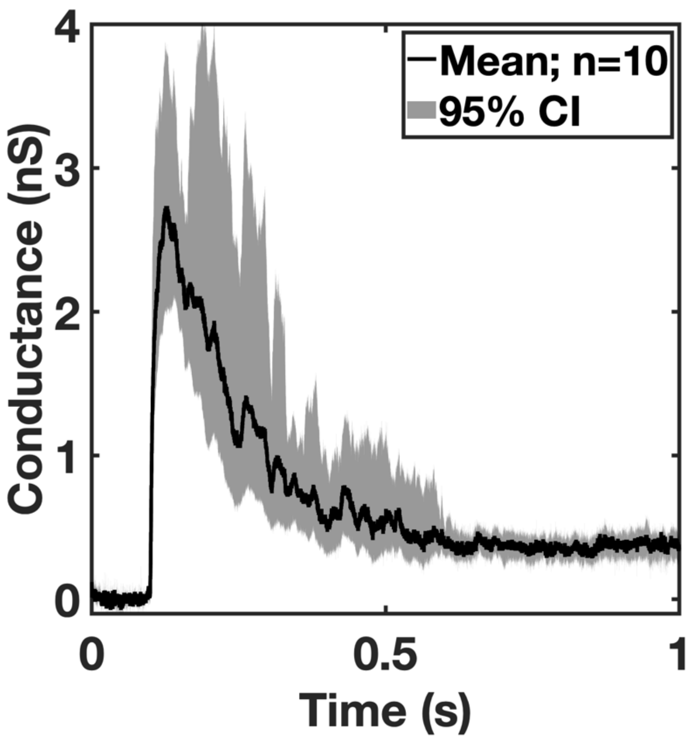


**Supplementary Figure 4.** Transients induced by WT parasites in low external calcium concentration (0.1 mM $CaCl_2$) have similar properties to WT transients induced in regular LCIS (2.0 mM $CaCl_2$; Figure 3a). The average waveform of low-calcium transients displays the characteristic fast rise to peak conductance and slower recovery to a new baseline (n=10). The confidence interval (CI) is calculated for each point along the averaged transient.

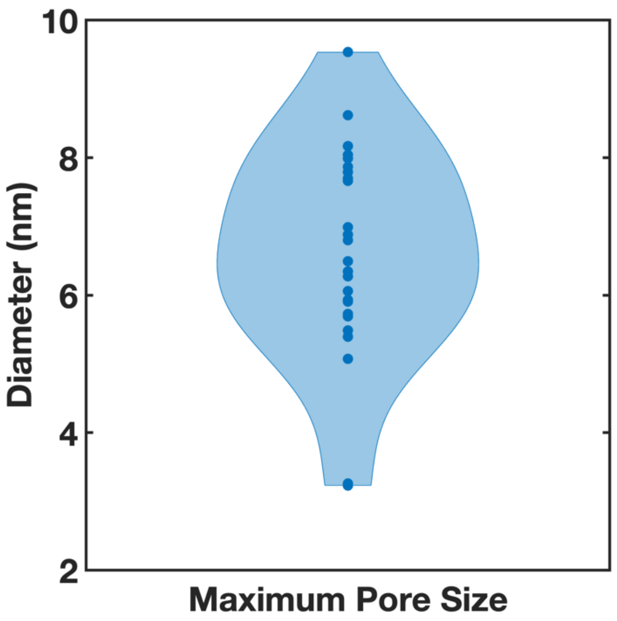


**Supplementary Figure 5.** Violin plot of pore diameters calculated from individual WT transient maximum conductance using a model for a cylindrical pore (see methods).
